# Modeling breeding programs considering social behavior in large groups of farmed fish

**DOI:** 10.64898/2026.02.05.704089

**Authors:** Gabriel Rovere, Beatriz C. D. Cuyabano, Florence Phocas

## Abstract

Breeding programs are essential in aquaculture, improving economically and environmentally important traits. In aquaculture systems, animals are raised in large groups, where social interactions are frequent and can influence individual performance. In these circumstances, indirect genetic effects can play an important role in the response to selection, and consequently, their effects on selection outcomes must be analyzed.

This study aimed to evaluate the implications of heterogeneous social interaction effects on fish breeding programs using stochastic simulations. We simulated a fish breeding program with 2000 selection candidates from 1000 families formed by a partial mating design of 100 males and 100 females. Social interactions were simulated, affected by the target phenotype and two latent-personality traits. We investigated how genetic gains and phenotypic variances are affected by the magnitude and direction of social interaction effects on the target phenotype, different selection strategies, and the genetic correlations between the target phenotype and personality traits. Our results showed that increased social interaction effects lead to greater phenotypic variability in the target trait. Under mass selection, the genetic means of personality traits change, and these changes depend on the strength and direction of genetic correlations between the focal and personality traits. Conversely, group selection did not increase phenotypic variability but reduced genetic gain for the focal trait compared to mass selection. Moreover, group selection did not alter the genetic means of personality traits. However, this approach increased the rate of inbreeding per generation, which could be mitigated by optimizing the number of families per group.

## INTRODUCTION

Selective breeding programs play a crucial role in aquaculture by improving traits of economic and environmental importance (Vandeputte *et al*., 2022; Pouil *et al*., 2025; De Montmorillon *et al*., 2026). In aquaculture, species are farmed in groups, ranging from hundreds to thousands, raised in cages, ponds, tanks, or raceways. Within these environments, social interactions are frequent and can strongly impact animal behavior and performance. When social interactions among individuals affect their performance, an individual’s phenotype reflects both its direct genetic effect (DGE) and the indirect genetic effects (IGEs) of its group mates. This means the environmental component of a phenotype can itself have a heritable basis (from group mates), implying that some part of the environmental component evolves along with the focal trait allowing it to evolve alongside the focal trait. Consequently, an individual’s genetic value affects not only its own phenotype, through individual selection, but may also affect the phenotype of group mates through social or associative effects.

The notion of associative effects and their consideration within selection theory was early presented in a series of contributions by (Griffing, 1967, 1968, 1969). Hence, the IGE have been modeled and integrated into the classical quantitative genetic models based on the variance-component model presented by (Griffing, 1967) and further developed by several authors (Muir, 2005; Bijma, Muir, and Van Arendonk, 2007; Bijma, Muir, Ellen, *et al*., 2007). In the field of evolutionary biology, Moore *et al*. (1997) introduced the concept of interacting phenotypes, and provided a model of the phenotypic evolution of traits affected by interactions among unrelated individuals. Based on previous contributions by Kirkpatrick and Lande (1989), the authors developed a general model that included the environmental effect on the focal individual’s phenotype, whether the group mates are related or not. The mathematical model divides the environmental effect into two components: the general environmental effect and the environmental effect due to the phenotype(s) of the group mate. Thus, the final observed phenotype of the focal individual has a component that is a direct function of the phenotypic trait(s) values of its social counterparts. This model, a trait-based model since the effector trait(s) of the group mate is(are) identified, includes an interaction coefficient ψ, a path coefficient that describes the extent to which the focal phenotype changes as a result of interaction with a group mate (Moore *et al*., 1997).

Following these initial models to account for IGE, more recent contributions have further developed the theoretical framework and quantitative modeling to better describe how IGEs affect phenotypes (Ellen *et al*., 2014; Bijma, 2014; Marjanovic *et al*., 2018). A modelling aspect addressed by these recent studies is the heterogeneity of IGE among individuals, an issue overlooked in early works, which assumed uniform, pairwise and homogeneous interactions. In aquaculture, where animals are reared in large groups, the assumption of homogeneous effects rarely applies. Instead, the group structure suggests that the frequency of interactions and their effects may differ considerably among individuals (Sloman *et al*., 2000; Luan *et al*., 2015; Backström *et al*., 2021).

The magnitude of social interactions (SI) on traits like growth, feed efficiency, and survival is well-documented in terrestrial livestock species (Peeters *et al*., 2012; Ellen *et al*., 2014; Alemu *et al*., 2014; Camerlink *et al*., 2014; Canario *et al*., 2017). While SIs are also recognized in fish, studies quantifying the magnitude of IGEs on economically relevant traits are limited (Brown and Brown, 1993; Monsen *et al*., 2010; Nielsen *et al*., 2014; Luan *et al*., 2015; Khaw *et al*., 2016). Underlying these SI are personality and behavioral traits, such as boldness, aggressiveness, sociability, explorative or cooperative behaviors, which shape group dynamics, and influence growth, stress, and overall performance of animals raised in groups (Camerlink *et al*., 2018; Angarita *et al*., 2019). These traits have been widely described in fish species, as they might influence survival, foraging efficiency, and dominance hierarchies (Sloman *et al*., 2000; Cañon Jones *et al*., 2010; Millot *et al*., 2014; Lallias *et al*., 2017; Best *et al*., 2023). Social interactions influenced by personality traits can modulate IGEs, either by amplifying positive behaviors such as cooperative feeding, or by exacerbating negative effects, such as competition and stress-induced growth suppression. Some studies indicate that fish may cluster in groups based on individuals that engage in more frequent and intense social behaviors, regardless of their size relative to others in the group (Sloman *et al*., 2000; Cañon Jones *et al*., 2010; Backström *et al*., 2021; Best *et al*., 2023).

Radersma (2021) proposed a framework using social networks to estimate genetic variation in social phenotypes, introducing two latent traits: social tendency (an individual’s willingness to interact) and social governance (its contribution to interactions). Wang *et al*. (2023) applied Radersma’s (2021) analytical framework to social behavior in poultry, defining two directional SIs roles, performer and receiver. According to these authors, an individual can play one of these two roles, being either the performer or the receiver of an action, and associate these two social tendencies with the indirect and direct genetic effects of the quantitative model presented by Bijma (2014).

Because of the influential role that IGEs might have on the rate and direction of response to selection, it is relevant to analyze their relationship with the DGEs to understand the expected response to selection (Bijma, 2010). Thus, this paper aimed to evaluate the impacts of different selection scenarios with heterogeneous IGEs on fish breeding programs through simulations. Specifically, we applied a model inspired in previous studies (Marjanovic *et al*., 2018; Radersma, 2021; Wang *et al*., 2023) on a fish breeding program with a partial factorial mating design (Dupont-Nivet *et al*., 2006; Vandeputte *et al*., 2022). Different selection scenarios were simulated for animals raised in large groups, where personality traits influenced the probability and intensity of animals’ interactions. These simulated scenarios were used to investigate the effects on genetic gains and variances when IGEs are present, according to: a) the magnitude and direction of the effects of the SIs on the target phenotype; b) the different selection strategies; and c) the direction of genetic correlations between the target phenotype and the personality traits that affect the SI.

## METHODS

### Model

A trait-based model was used to define the social partners’ indirect genetic effects on the phenotype of a focal individual. Assuming that the same trait is at play in both focal and interacting individuals, the model for a target trait in the focal individual i can then be expressed as follows (Moore *et al*., 1997):

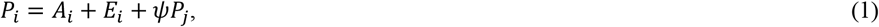

where P_i_ is the phenotype of the focal individual *i*, which, in a classical quantitative model, is partitioned in the additive genetic effects (A_i_) and the environmental effects. (Moore *et al*., 1997) model, however, separates the environmental effects in two components. The first component is E_i_, an environmental effect independent of the social environment of individual *i*. The second component is *ψ*P_j_, which represents the effect of the phenotype of social partner *j* (P_j_) on the phenotype of individual *i* (P_i_); *ψ* is a coefficient that expresses the degree to which the phenotype of individual *i* is affected by its interaction with individual *j*.

Based on the model described by equation (1), Marjanovic *et al*. (2018) proposed a quantitative genetic model involving the interaction of two individuals using body weight as an effector trait in aquaculture. The authors proposed that the effect of SIs on the phenotype of the focal individual was b_ij_(P_j_-P_i_), in a model similar to the following,

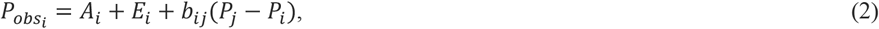

Where 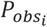 is the phenotype observed for body weight on the focal individual after the SI, b_ij_ is a regression coefficient that resembles *ψ* in equation (1), and P_i_ and P_j_ are the phenotypes for body weight of focal individual *i* and its social interactor *j*.

Marjanovic *et al*. (2018) proposed *b*_*ij*_ as a specific parameter for every interacting couple, defined as *b*_*ij*_= *b* + A_Di_ + E_Di_ + A_Ij_ + E_Ij_, where b represents the average regression coefficient, considered as a population parameter indicating whether SIs are predominantly competitive (*b*<0) or collaborative (*b*>0). The other terms of the equation that defines *b*_*ij*_ are the direct genetic and environmental effects (A_Di_ and E_Di_ respectively) of the focal individual i, and the indirect genetic and environmental effects (A_Ij_ and E_Ij_ respectively) from the individual j. In this configuration, the model considers the trait that accounts (effector trait) for the SI is the same in both interactor individuals, and allows asymmetrical SIs. Our present study aimed to incorporate the ideas proposed by Radersma (2021) and Wang *et al*. (2023) in a quantitative model that considers personality traits, as effector traits on the social environment in which an individual *i* interacts with a social partner *j*. Our model was based on that presented by Marjanovic *et al*. (2018), and is described by equation (2), with b_ij_ modified to be determined by personality traits (Radersma, 2021; Wang *et al*., 2023).

We defined two personality traits related to behavior tendency, one describing the behavior on the axis of aggressiveness (trait P), and the second describing the behavior on the axis of stress response (trait R). A positive value of trait P (i.e. a more aggressive behavior) will incur a negative impact on a more sensitive interactor (i.e. with a positive value of R) to environmental stressors (Gesto, 2019). Traits P and R were defined to resemble the terminology used by Wang *et al*. (2023) where P described the tendency of an individual to perform an action towards others, and R described the tendency of an individual to be the receptor of an action from others. Finally, the effect of the difference of two individuals’ body weights on the observed phenotype 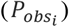 is implicitly weighted by their personality traits, which determine the strength of the coefficient *b*_*ij*_ in equation (2), as follows:

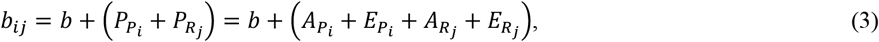

where *b* is a population parameter that indicates the predominant type of SIs in the population, being negative under a predominantly competitive environment, and positive under a predominantly cooperative behavior of the interactors; 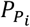 and 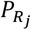, are the phenotypes of the personality traits P and R previously described, in the focal individual *i* and its social interactor *j* respectively. 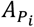 and 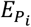, are the corresponding additive genetic and environmental values of trait P of the focal individual i, and 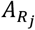 and 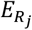 are the corresponding additive genetic and environmental values of trait R of the social interactor j.

### Simulation

We used simulated data to assess the effect of different selection methods under different social environments (competitive, neutral, or collaborative), on the mean and variance of a phenotype such as the body weight in fish, that in this context operates as a social and effector phenotype. The breeding scheme was based on a partial-factorial mating plan, commonly used in French fish commercial breeding programs (Dupont-Nivet *et al*., 2006; Vandeputte *et al*., 2022). The breeding scheme had a nucleus of 200 parents (100 males and 100 females) mated in groups with equal proportions of sex and with a phenotyping constraint of 2,000 individuals. We started by simulating 200 unrelated founder individuals to generate 2,000 offspring that defined the base population, from which we randomly chose 200 parents to finally start 10 generations of phenotypic selection under different group selection strategies.

### Founders and Base Population

Two hundred unrelated founder individuals were simulated with three traits: one trait was the phenotype of interest without the effect of SI (P), and the other two were personality traits with an incidence on the individual’s behavior in SIs (P_P_ and P_R_).

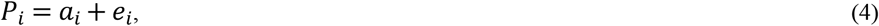

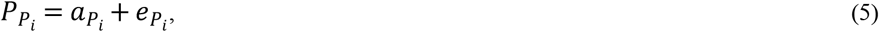

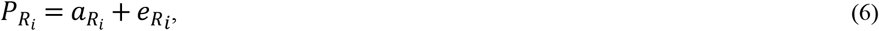

The additive genetic effects were sampled from a multivariate normal distribution 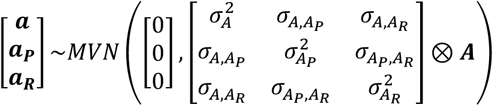, where **A** is the numerator relationship matrix, 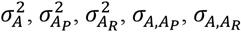 and 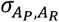 are the additive genetic variances and covariances between the three traits. Similarly, the environmental effects (E) were sampled from multivariate normal distribution 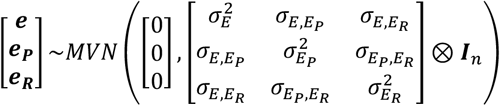, where ***I***_*n*_ is the identity matrix, 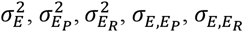 and 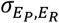 are the environmental variances and covariances between the three traits.

In the founder population, the three traits were uncorrelated, and for all of them phenotypic variance was assumed to be *σ*^2^ = 3, the genetic variance was 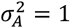 and the residual variance was 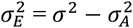. For all three traits, the breeding values (i.e. the additive genetic effects) of the offspring in subsequent generations were obtained from parents’ average plus a Mendelian sampling term, i.e. as 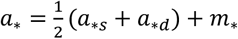, where s and d correspond to the sire and dam’s additive genetic effects, the ‘∗’ notation generalizes this breeding values for either the trait of interest or the personality traits, and *m*_*_ corresponds to the mendelian sampling term, which was sampled from a multivariate normal distribution for the three traits.

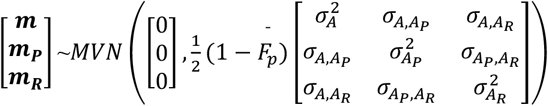, where 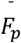 is the average inbreeding coefficient of the parents.

### Social interactions

We simulated SIs for the base population and subsequent generations, modelling two components: the probability of interaction between two individuals, and the outcome of such SI, if it occurred. Following previous studies (Huntingford *et al*., 2012; Marjanovic *et al*., 2018, 2022), we considered that the relative size of fish can have an incidence in their interactions. SI probability was further modulated by individual personalities and sizes. When individuals *i* and *j* meet in the tank, the probability (*π*_*ij*_) that they interact depends on how much individual i expresses a competitor (or collaborative) behavior (P_Pi_) and how much individual j is resistant (or susceptible) to interact (P_Rj_), and on their relative sizes (P_i_, P_j_). Since interactions are bidirectional, *π*_*ij*_ splits equally: half reflects i imposition on j, and half on the reverse, as described by equation (7):

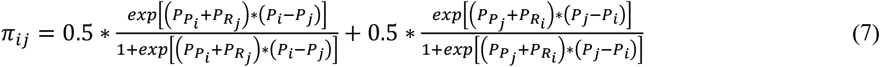

An interaction between individuals can only occur if they meet. Thus, the final probability of individuals i and j interacting was *π*_*ij*_, defined by their sizes and personalities, weighted by their spatial proximity in the tank. To model physical proximity we assigned each individual three random coordinates (*x*_1_, *x*_2_, *x*_3_) and calculated the pairwise Euclidean distance between group mates. Without any loss of generality, we scaled the Euclidean distance by the maximum distance, ensuring independence from fish density (fish/m^3^) and thus,

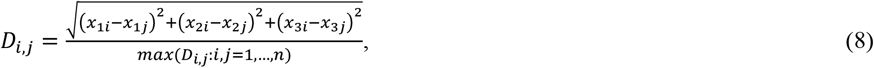

where *D*_*i,j*_ is the scaled Euclidian distance between individuals *i* and *j*. The individual’s proximity is then given by (1 − *D*_*i,j*_), and finally, the probability of interaction considering the physical proximity between interactors was:

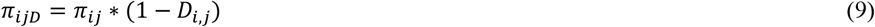

In the simulations, we used Bernoulli sampling with probabilities *π*_*ijD*_ for each pair of individuals (i, j) to determine if an interaction occurred,

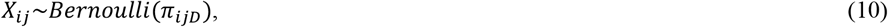

such that *X*_*ij*_ = 1 indicated that the interaction occurred, and *X*_*ij*_ = 0 indicated that it did not take place.

Thereafter the result of the SI between *i* and *j* was given by:

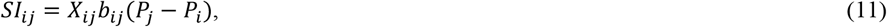

where *b*_*ij*_ is the regression coefficient between *i* and *j*, as defined in equation (3), and *P*_*i*_ and *P*_*j*_ are their respective phenotypes for the focal trait (size), before the SI. Finally, the observed phenotype of the focal individual *i* was the sum of its baseline phenotype and the results of all its SIs weighted by a scaling factor *γ*, which will be explained shortly,

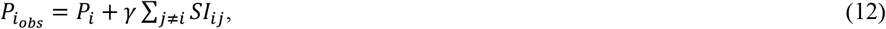

where *γ* is a scaling factor to ensure that the SIs do not add an unrealistic amount of variance to the observed phenotype in equation (12), and to allow a control in the simulation routine, of how much variance is due to the SIs at the base population. The variance of the observed phenotype in equation (12) is given by 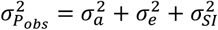, such that 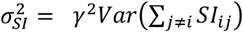. Therefore,*γ* can be defined as:

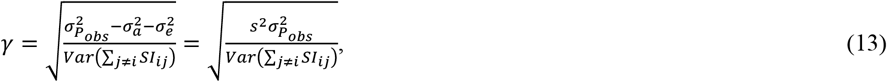

where *s*^2^ is the portion of variance of the observed phenotype in equation (12), explained by the SI.

While the values of 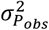 and *s*^2^ can be defined directly as parameters, obtaining an analytical value for *Var*(Σ_*j*≠*i*_ *SI*_*ij*_) is not straightforward. That is because each *SI*_*ij*_ depends on the Bernoulli *X*_*ij*_, with probabilities given as non-linear functions of three phenotypes inherent to the individuals, and then the final *SI*_*ij*_ is a multiplication between *X*_*ij*_, *b*_*ij*_ (dependent on the personality traits), and the difference between the individuals’ sizes (*P*_*j*_ − *P*_*i*_). Despite the challenge of analytically deriving *Var*(Σ_*j*≠*i*_ *SI*_*ij*_), in a simulation setup, the scaling factor *γ* can be empirically defined at the base population by simply replacing *Var*(Σ_*j*≠*i*_ *SI*_*ij*_) in equation (13) with its observed value 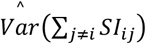.

For each simulation replicate, given that both 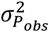 and *s*^2^ were defined, the scaling factor *γ* was calculated empirically following equation (13), and its value was kept fixed for every subsequent generation.

### Scenarios simulated

All scenarios began with a nucleus of 200 parents (100 males and 100 females), mated in 10 groups of 10 males and 10 females each. This produced 100 families per group (1,000 families total), with 200 offspring per group (2 full-sibs per family), using the parameters described in the previous section. The 2,000 offspring were raised together, with SIs simulated to form the base population. From this population, 200 individuals were randomly selected as parents for generation 1.

The base scenario assumed SIs had no influence on the focal trait’s phenotype, and mass selection was applied based on observed phenotypes of the focal trait. The focal and personality traits were assumed uncorrelated, with variance components as described before. Alternative scenarios explored the effects of:

a. Magnitude of SI effects: We compared the base scenario (0% variance explained by SIs) to scenarios where SIs explained 10% and 20% of the phenotypic variance in the focal trait.
b. Direction of SI effects: We tested three scenarios for *b*: competitive (*b* = -0.5), neutral (*b* = 0), and collaborative (*b* = +0.5).
c. Selection strategy: In the scenario where SIs explained 10% of the phenotypic variance, we compared mass selection to group selection using two family structures: progeny groups from either 10×10 or 5×5 parent combinations of each sex.
d. Correlation between personality traits and the focal trait: We examined three genetic correlation scenarios between the focal trait and the two personality traits (with SIs explaining 10% of phenotypic variance): no correlation (0), negative correlation (-0.5), and positive correlation (+0.5). Environmental correlations were assumed equal to genetic correlations, and personality traits were uncorrelated.

The base scenario was simulated using the same strategies as the others. It was designed to represent a situation where SIs explained 0%—not the absence—of the phenotypic variance in the focal trait, allowing us to observe the evolution of social behavior under this condition. Consequently, results from the base scenario across different b values were expected to be statistically identical. The SI effects were simulated for the base population with *b* equal to zero, to generate the observed phenotype that later was used to start the following 10 generations (Fig. 1).We simulated 50 independent replications of each scenario, using a vector of integers to start the random number generator. We used Tukey’s honest significance test at a significance level of 0.05 to evaluate if the differences between scenarios were statistically significant at every generation of selection. The fixed model used for the analysis of variance took into account the following factors: the effect of repetitions (r = 50 levels), the effect of scenarios (s = the number of levels depending on the number of scenarios compared), the effect of *b* (b = 3 levels), and the interaction between scenarios and b (with the number of levels being s * b), and a random residual term. The simulations were performed using self-made code in R (R Core Team, 2025).

**Fig. 1.**
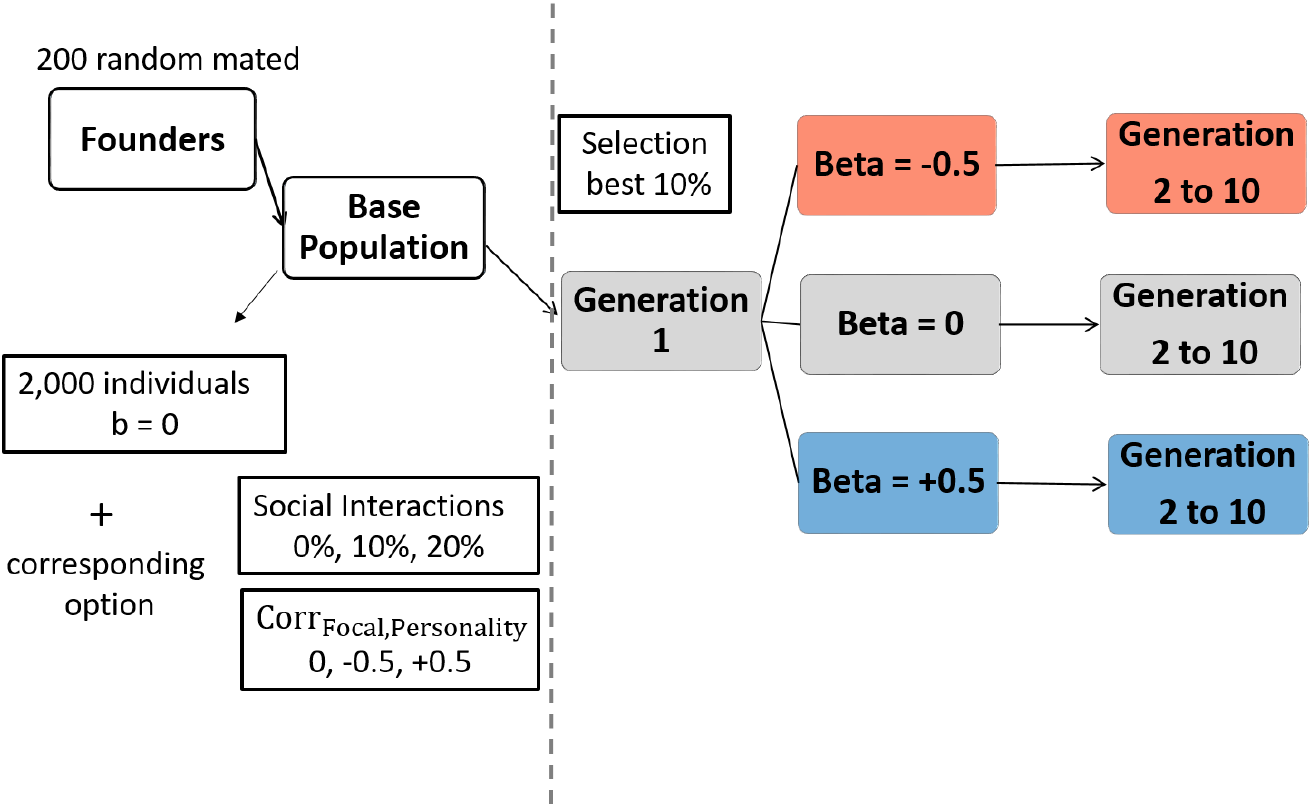
Diagram to illustrate the simulation scheme.

## RESULTS

### Effect of the magnitude and direction of the Social Interaction effects

Figure 2 illustrates how the current mating system (10×10 parents) and mass selection influence the evolution of the observed phenotype across three scenarios of SI magnitude. In Fig. 2.A., the focal phenotype means at generation 10 were highest in scenario with highest SI effects (Table S1.a). Specifically, in SI20 scenarios, where SIs accounted for 20% of the phenotypic variance of the focal trait, the phenotypic mean was significantly higher under competitive SI environment (*b*= -0.5) compared to neutral (*b* = 0) or collaborative (*b* = +0.5) ones. For the SI10 scenarios, where SIs accounted for 10% of the phenotypic variance, neither the neutral nor the collaborative SI environments showed significant differences from the base scenario (no SI effect) (Table S1.a).

**Fig. 2.**
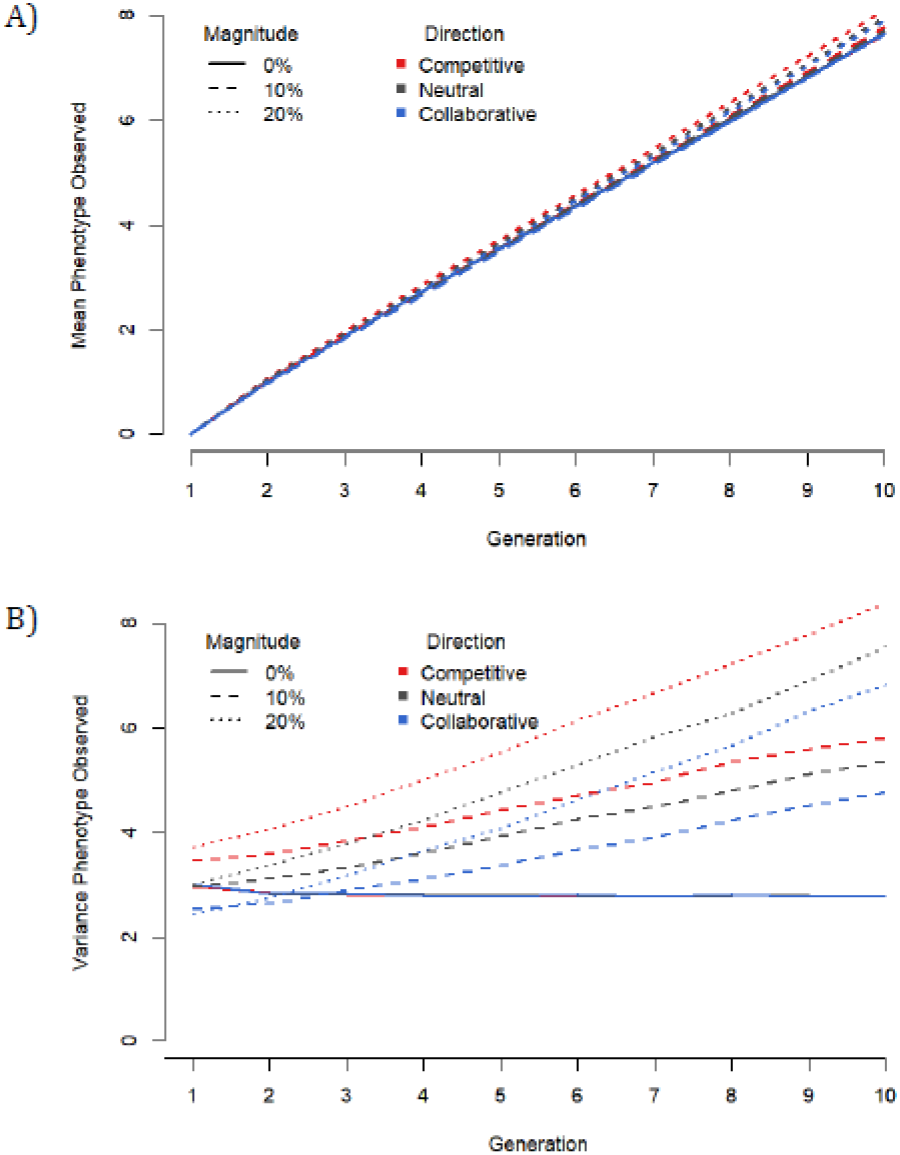
Effect of the different magnitude of social interactions’ effects on the focal trait along 10 generations of mass selection on: A) individual mean of phenotype observed; B) the variability of the phenotype observed.

However, the variance of the observed phenotype differed markedly across all scenarios where SIs influenced the focal phenotype (Fig. 2.B.). Over the generations simulated, phenotypic variance increased in every SI scenario, with the highest variance in competitive SIs (*b* = -0.5), and the lowest in collaborative ones (*b* = +0.5). When SIs had no effect on the observed phenotype, phenotypic variance instead declined over the 10 generations, with no differences observed between *b* scenarios, as expected (Fig. 2.B, Table S1.b).

Considering the original parameters of the base population for the observed phenotype (*μ* = 0, *σ*^2^ = 3), the mean in the base scenario increased by ∼ 4.4 phenotypic standard deviations (*σ*) by generation 10. In SI20 scenarios, the phenotypic mean changed ∼ 4.7*σ*. At the genetic level, the SI20 competitive scenario had the highest mean at generation 10, while the SI10 collaborative scenario had the lowest; the remaining scenarios showed statistically similar genetic means (Fig. 3.A and Table S2.a). The personality trait P, associated with willingness of to engage in social activity, declined in genetic mean over generations. This decrease was most pronounced in the SI20 scenario with *b* = +0.5, though the change was less marked than the genetic change observed in the focal trait (Fig. 3.B, Table S2.b).

**Fig. 3.**
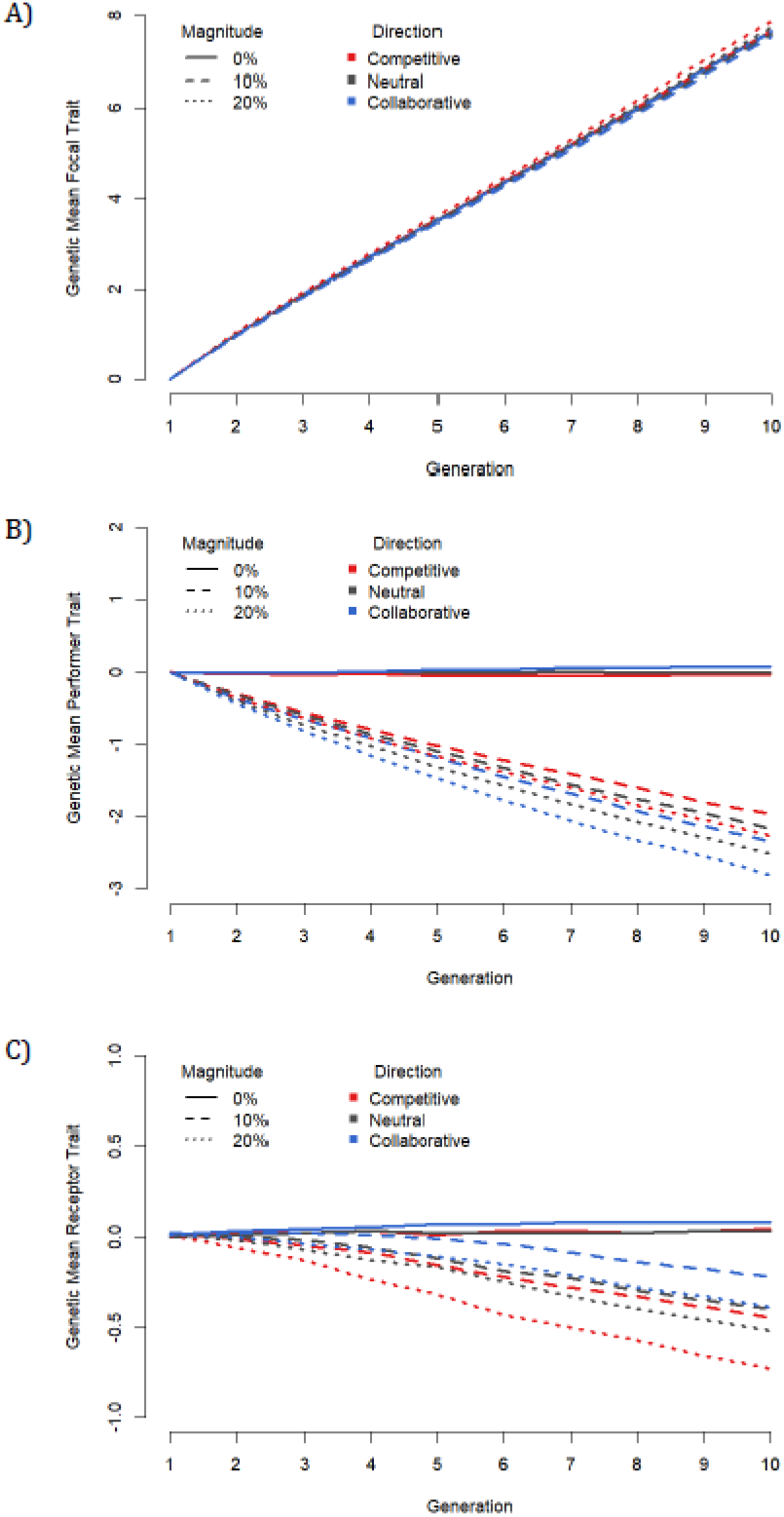
Effect of the different magnitude of social interactions’ effects on the focal trait along 10 generations of mass selection on: A) the mean breeding value of focal trait; B) the mean breeding value of the personality trait, “perfomer”, linked to axis competitive-collaborative; C) the mean of the breeding value of the personality trait, “receptor”, linked to stress or resilience.

Similarly, trait R, associated with susceptibility to SIs, also showed a negative genetic response, but the decrease was smaller than that observed for trait P (Fig. 3.B, Table S2.c).

### Effect of different selection strategies

Assuming SIs account for 10% of the phenotypic variance in the focal trait, we evaluated three selection strategies:

a. Mass selection (MASS), where the top 10% of candidates were selected as parents for the next generation;
b. Group selection (GR10), using 10 groups formed by factorial mating plans of 10 sires and 10 dams; the best-performing group provided the parents for the next generation;
c. Group selection (GR20), using 20 groups formed by factorial mating plans of 5 sires and 5 dams; the two best-performing groups provided parents for the next generation.

Figure 4.A shows the evolution of the observed phenotype mean over 10 generations. Although group selection (GR10 and GR20) initially seemed to outperformed mass selection (MASS), by generation 10, MASS had the highest mean, although GR20 still performed significantly better than GR10. mean social environments (*b*) showed significant differences (Fig. 4.A. and Supplemental Table S.3.a). However, phenotypic variability strongly differed between selection strategies (Fig. 4.C).

**Fig. 4.**
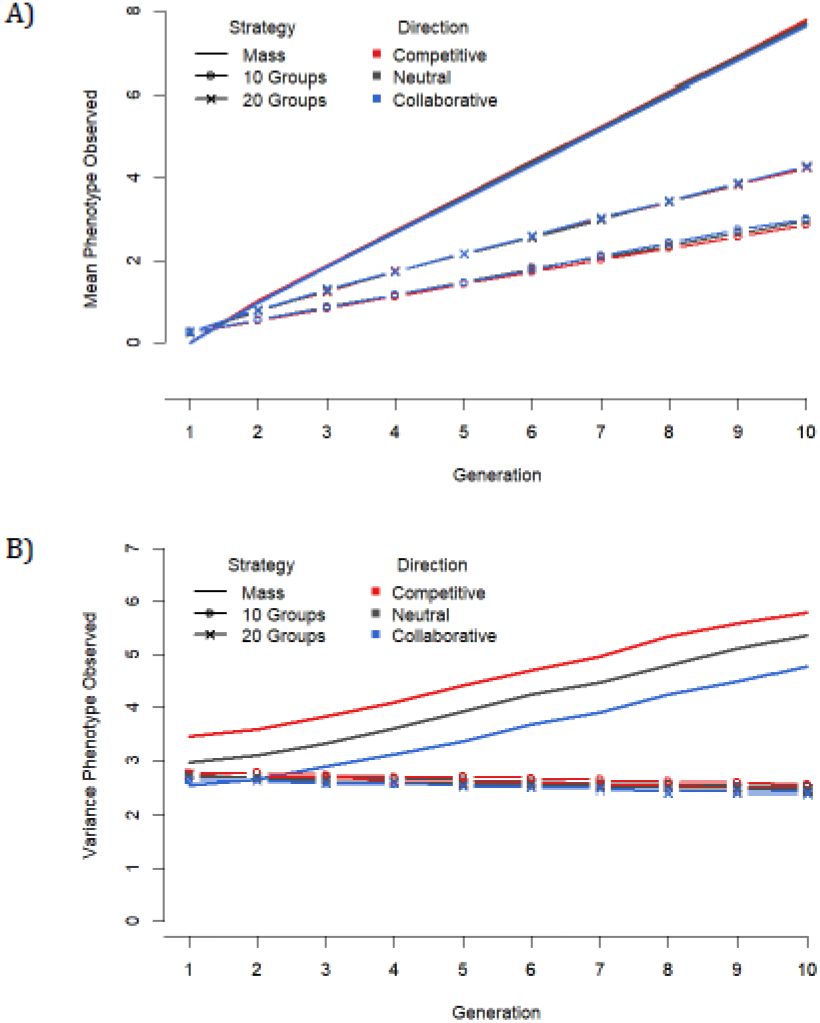
Effect of three selection strategies along 10 generations, with a scenario of 10 % of the phenotypic variance explained by social interaction effects of the focal trait, on: A) individual mean of phenotype observed; B) the variability of the phenotype observed. Mass = mass selection, 10 Groups= group selection from 10 groups, 20 Groups = group selection from 20 groups.

Under mass selection, variance in the observed phenotype increased continuously, especially in competitive SI environments, whereas under both group selection strategies, phenotypic variance remained stable or slightly decreased (Fig. 4.C, Table S3.b). It is important to note that while GR20 achieved significantly higher phenotypic improvement than GR10, it did not exhibit the same concomitant increase in variance observed in the MASS scenario. In summary, at the phenotypic level, mass selection increased both the phenotypic mean and variance to values 1.8 times higher than those achieved by the best group selection strategy (GR20). Group selection based on 20 groups (GR20) improved phenotypic results by approximately 50% over GR10 after 10 generations, without increasing phenotypic variance. Similar results were obtained for the genetic trend of the focal trait (Fig. 5.A.). By generation 10, the genetic mean under mass selection was approximately 1.9 times higher than with GR20, and GR20 mean was approximately 1.5 times higher than GR10 one. No significant differences were observed between different directions of SI effects (Fig. 5.A, Table S4.a).

**Fig. 5.**
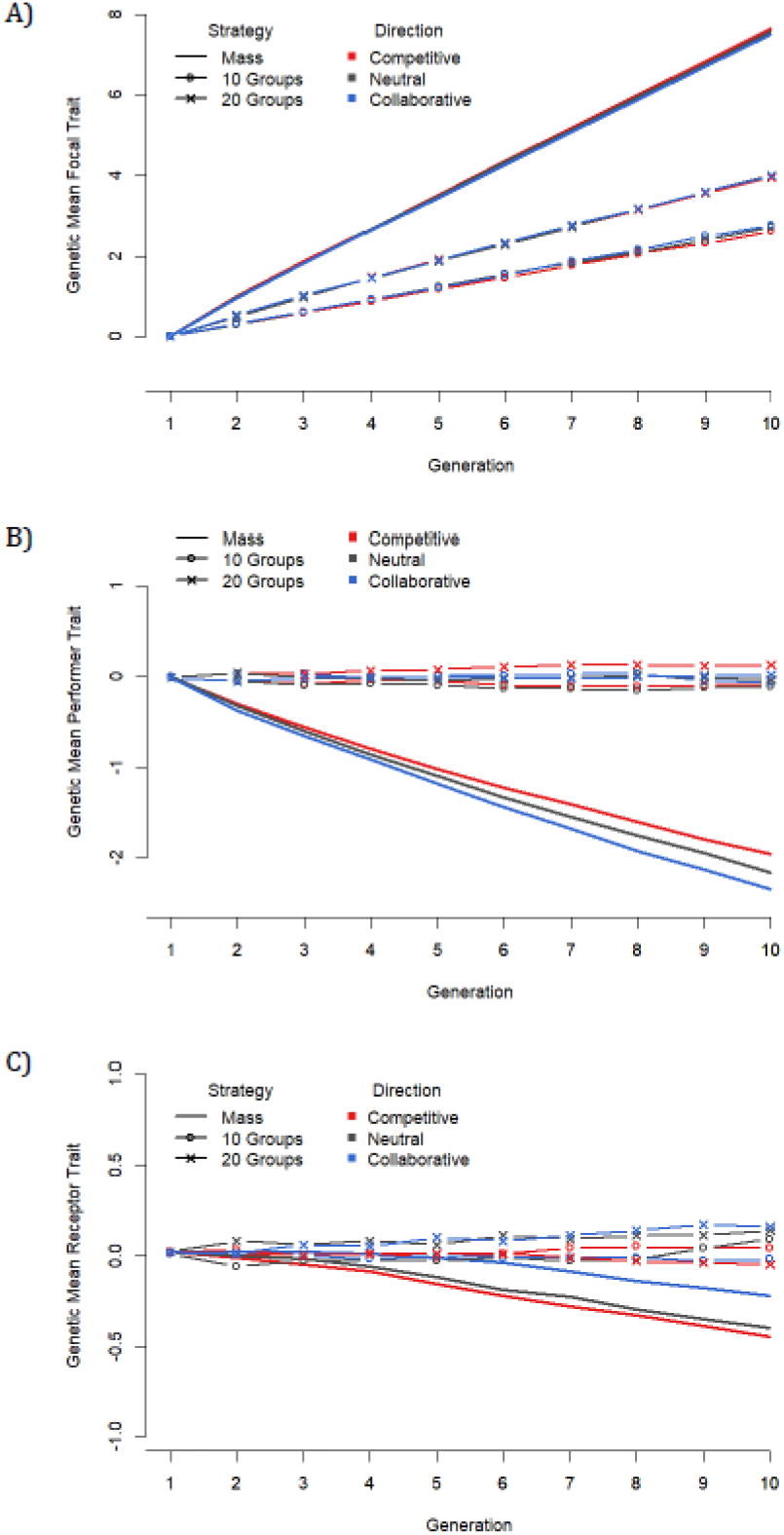
Effect of different selection strategies along 10 generations, with a scenario of 10 % of the phenotypic variance explained by social interaction effects of the focal trait, on: A) the mean breeding value of focal trait; B) the mean breeding value of the personality trait, *“* performer*”*, linked to axis competitive-collaborative; C) the mean of the breeding value of the personality trait, *“*receptor*”*, linked to stress or resilience. Mass = mass selection, 10 Groups= group selection from 10 groups, 20 Groups = group selection from 20 groups.

However, it is important to underline that while the mean inbreeding level increased moderately under mass selection (∼ 0.05 at generation 10), it was strongly increased under group selection with a mean ∼0.22 by generation 10. Between group selection strategies, GR20 showed a slightly but significantly higher mean inbreeding level than GR10 by generation 10 (Table S4.d).

For the personality traits, the most substantial genetic changes occurred under mass selection, particularly for trait P. While the genetic mean of trait P decreased in MASS scenario, especially in collaborative SI environment (Fig. 5.B, Table S4.b), this change was negligible under group selection. No significant differences were found between GR10 and GR20, nor between different directions of SI effects.

For trait R, differences between selection strategies were minor, though some were statistically significant. Under group selection, trait R showed either no genetic change or a slight positive trend when SI effects were neutral or collaborative (*b* = 0 and *b* = +0.5). In contrast, mass selection led to a slight negative genetic trend, particularly in competitive SI environments (*b* = -0.5) (Fig. 5.C, Table S4.c).

### Effect of the correlation between focal trait and personality traits

Assuming a scenario where SI effects account for 10% of the phenotypic variance in the focal trait, we simulated three scenarios of genetic and environmental correlation between the focal trait and the two personality traits: uncorrelated (0), negatively correlated (-0.5), and positively correlated (+0.5). In each scenario, genetic and environmental correlations were assigned the same value.

Figure 6.A presents the results for the observed phenotype of the focal trait. By generation 10, the mean observed phenotype was highest when correlations between the focal and personality traits were negative, compared to the uncorrelated scenario. The lowest mean occurred when correlations were positive. Starting from generation 5, scenarios with negative correlations showed a significantly higher phenotypic mean than the other two correlation scenarios. Differences between the uncorrelated and positively correlated scenarios became significant from generation 7 onward.

**Fig. 6.**
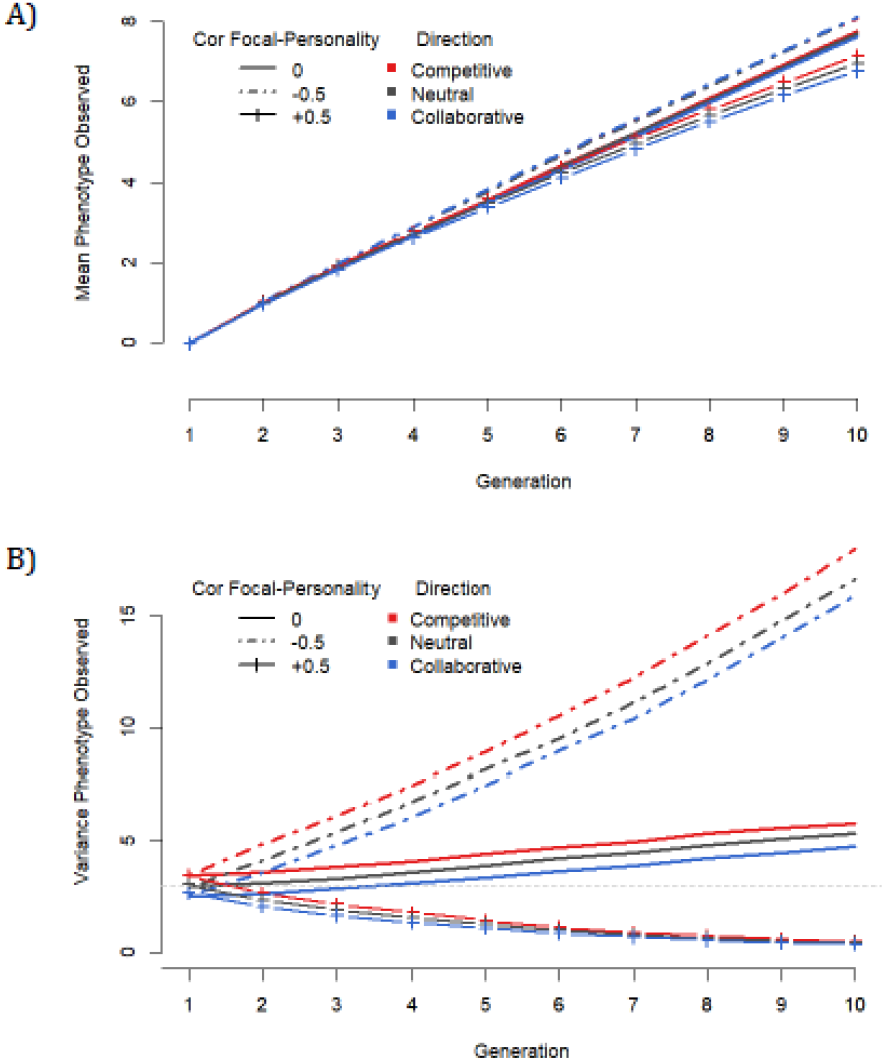
Effect of the direction of the genetic and environmental correlation between focal and personality traits with a scenario of 10 % of the phenotypic variance of the focal trait affected by social interaction effects and mass selection on: A) individual mean of phenotype observed; B) the variability of the phenotype observed. 0 = Genetic and environmental correlation between Focal trait and Personality traits is zero; -0.5 = Genetic and environmental correlation between Focal trait and Personality traits is -0.5; +0.5 = Genetic and environmental correlation between Focal trait and Personality traits is 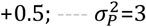.

By generation 10, the direction of SI effects made a significant difference only in scenarios with positive correlations. In these cases, the highest mean values occurred when *b* was negative, while the lowest values occurred when *b* was positive. As a result, the difference in mean phenotype at generation 10 varied by approximately 0.8*σ* between the two most contrasting scenarios: the scenario with negative correlations between focal and latent traits (∼4.7σ) and the scenario with positive correlations and collaborative SI effects (∼3.9σ) (Fig. 6.A, Table S5.a).

The scenario with negative correlations between focal and latent traits also exhibited the highest phenotypic variance. This increase in variance became more pronounced as the environment shifted from collaborative (*b* = +0.5) to competitive (*b* = -0.5). When traits were uncorrelated, variance increased less, particularly when SI effects were collaborative (*b* = +0.5). Contrarily, in scenarios with positive correlations between focal and latent traits, the phenotypic variance of the focal trait strongly reduced by generation 10, with no significant differences observed between competitive, neutral, and collaborative environments.

Figure 7 presents the genetic changes in the focal and latent traits over 10 generations, considering their correlations. The focal trait exhibited a significantly higher genetic mean at generation 10 when was negatively correlated with the latent traits. When the focal trait was positively correlated with the personality traits, its genetic mean at generation 10 was approximately 1*σ* lower than in the other two correlation scenarios. In this case of positive correlation, the genetic mean of the focal trait was highest under competitive SI (*b* = -0.5) and lowest under collaborative SI (*b* = +0.5), with a significant difference of approximately 0.22*σ* between these two contrasting scenarios (Fig. 7.A, Table S6.a).

**Fig. 7.**
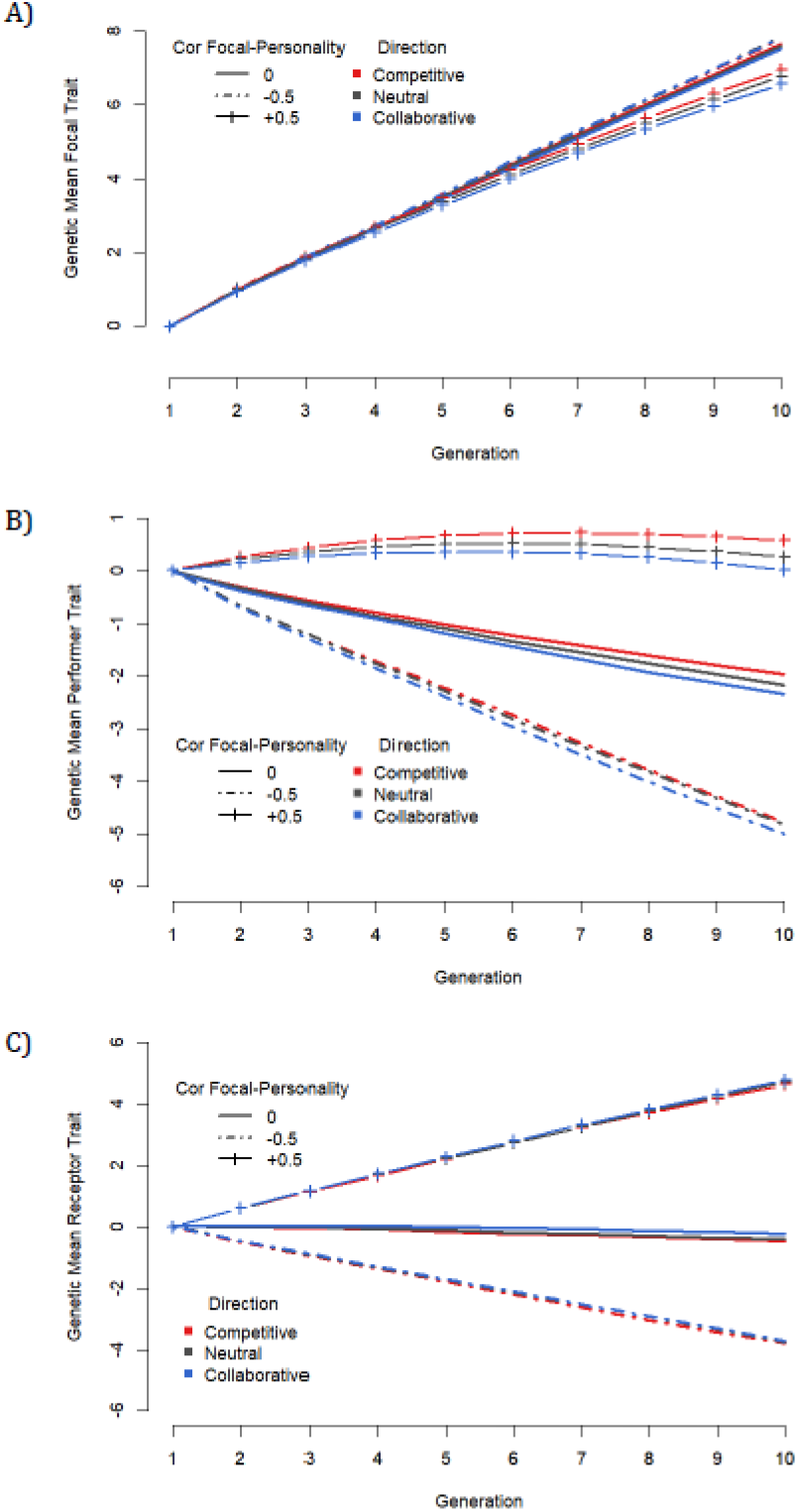
Effect of the direction of the genetic and environmental correlation between focal and personality traits with a scenario of 10 % of the phenotypic variance of the focal trait affected by social interaction effects and mass selection on: A) the mean breeding value of focal trait; B) the mean breeding value of the personality trait linked to axis competitive-collaborative; C) the mean of the breeding value of the personality trait linked to stress or resilience. 0 = Genetic and environmental correlation between Focal trait and Personality traits is zero; -0.5 = Genetic and environmental correlation between Focal trait and Personality traits is -0.5; +0.5 = Genetic and environmental correlation between Focal trait and Personality traits is +0.5.

The genetic trends of the personality traits varied distinctly across the three correlation scenarios. For trait P, a slight positive genetic trend was observed when the focal and personality traits were positively correlated. This trend turned negative when traits were uncorrelated and became even more negative when the correlations were negative. In all cases, the genetic trend for trait P was highest under competitive SI (*b* = -0.5) and lowest under collaborative SI (*b* = +0.5) (Fig. 7.B, Table S6.b). Trait R followed a similar pattern of genetic changes at generation 10, although the changes were more pronounced than for trait P in scenarios with positive correlations with the focal trait. On the contrary, in uncorrelated or negatively correlated scenarios, the decrease in the genetic mean of trait R was more moderate compared to trait P. The direction of SI effects did not influence the direction or magnitude of the genetic trend for trait R, except in the uncorrelated scenario, where the greatest decline in genetic mean occurred under competitive SI and the smallest decline under collaborative SI (Fig. 7.C, Table S6.c).

## DISCUSSION

In this study, we aimed to explore the expected evolution of phenotypic and genetic means and variances under phenotypic selection for a focal trait influenced by SIs in fish breeding schemes. We considered the magnitude and direction of these interactions, as well as the selection strategy applied, either mass or group selection.

### Differences and similarities with previous simulation studies on social interaction traits

In general terms, our findings align with previous research showing that mass selection applied to a focal trait influenced by SIs leads to genetic improvement in that trait, alongside an increase in its phenotypic variance. Unlike previous studies on this topic in aquaculture (Marjanovic *et al*., 2018, 2022), our work explicitly simulated SIs within tank groups. We assumed that both the probability of SIs and their effects depended on the focal phenotype and two personality traits. Despite these assumptions, our overall results agreed with Marjanovic *et al*. (2018), indicating that the magnitude and direction of SI effects impact the phenotypic variance of the focal trait more strongly than its phenotypic or genetic gain. We also confirmed that the selection strategy significantly influences both outcomes.

As in previous studies by Wang *et al*. (2023) and Hansson *et al*. (2025), we simulated individuals in a tank with varying probabilities of interacting. In our study, these probabilities were a function of the focal trait and the two latent personality traits, weighted by the physical distance between the individuals. Instead of following longitudinal time interactions, we made a simplification. We considered a one-time point interaction and used the scaled sum of interactions at this point as a proxy for the lifetime SIs experienced by an individual, which we saw as a potential limitation of our study. In our simulation, the focal phenotype served as an effector trait, but its effect was modulated by the personality traits. The weighting coefficient b_ij_ represented the deviation from the mean general personality interactions defined by *b*. Thus, the absolute value of b_ij_ reflected the strength of the SI, analogous to the idea used in Radersma (2021), with asymmetric weights between individuals (b_ij_ ≠ b_ji_). With similar approach, Wang *et al*. (2023) estimated breeding values for behavioral (“personality”) traits, exploring their relevance to commercial breeding programs. They simulated animal movement to model the SIs, a more detailed approach than ours, which simplified interaction probabilities based on physical distance using random coordinates assigned to individuals. We scaled the distances to maintain a constant density of individuals per cubic meter. Then, the event that an interaction occurred was purely determined by probabilities and not spatial constraints.

Hansson *et al*. (2025) modelled random networks in dairy cows using an average number of individuals per network based on literature. They modelled the frequency of interactions between cows within a group, to evaluate the effect of the frequency in their model. In our study, we assumed that all individuals could potentially interact, with SIs frequency as a function of personality and focal traits, and physical proximity within the tank.

### Magnitude of SI

Under the assumption that the focal and personality traits were uncorrelated, three scenarios of magnitude of SI effects were simulated, 0%, 10% and 20% of the phenotypic variance. The results showed that the phenotypic mean of the focal trait tended to increase in scenarios with greater magnitude (SI20) and competitive SIs (*b*= -0.5). However, the advantage observed in scenarios with greater SI effects at the phenotypic level did not correspond to genetic differences for the focal trait. Except for scenario SI20 and competitive SIs, the genetic mean was lower than in the base scenario, indicating a loss in the efficiency of the phenotypic selection. Additionally, the variance of the focal phenotype increased proportionally to the magnitude of the SI effects, regardless of whether the SI effects were competitive or collaborative. Indeed, within a specific scenario of magnitude of the SI effect, the variance of the observed phenotype increased from collaborative to competitive interactions, in line with the results reported by Marjanovic *et al*. (2018). Suppose we associate the focal phenotype with body size or body weight. In that case, that might imply an increasing loss of uniformity of these phenotypes, an undesirable result for commercial farming systems. With focal and personality traits uncorrelated, the current mass selection strategy would achieve similar genetic response beside the magnitude of the influence of SI effects on the focal phenotype. However, it would produce individuals with a greater tendency to engage in competitive behavior and less resilience to SIs (Fig.3.A and 3.C), and might move the mean SI environment to more competitive (Fig. 8.A). Studies on real populations under selection in aquaculture are limited, and despite all recognized the importance of the IGE, they reported a wide range of results in terms of the amount and direction of the SI on the phenotypes observed (Nielsen *et al*., 2014; Luan *et al*., 2015; Khaw *et al*., 2016). Khaw *et al*. (2016) concluded that in Nile Tilapia traditional selection, in an environment where the fish have to compete with each other for the resources, would increase competition, with a strong effect on harvest weight. In the Atlantic cod, Nielsen *et al*. (2014) did not find differences in growth traits between models with or without IGE, but significant contribution for length of the first dorsal fin and fin erosion, traits that might be welfare indicators. Luan *et al*. (2015) reported in Pacific white shrimps that SIs contributed to a large part of the heritable variation in adult body weight with non-significant positive direct–indirect genetic correlation. The authors interpreted this weak correlation as a neutral or slightly cooperative heritable interactions, rather than competition; however, they underlined that it may be due to the low rearing density. Studies at smaller scale, with experimental populations suggests that individuals with a competitive-aggressive profile might have higher stress levels. This increased stress could result in lower feed conversion and weight gain, or increased disease susceptibility (Cañon Jones *et al*., 2010; Lallias *et al*., 2017; Gesto, 2019; Best *et al*., 2023). Additionally, such profiles could lead to social networks characterized by dominance and subordination, leading to increasingly aggressive behavior, which could impact production efficiency (Backström *et al*., 2021).

**Fig. 8.**
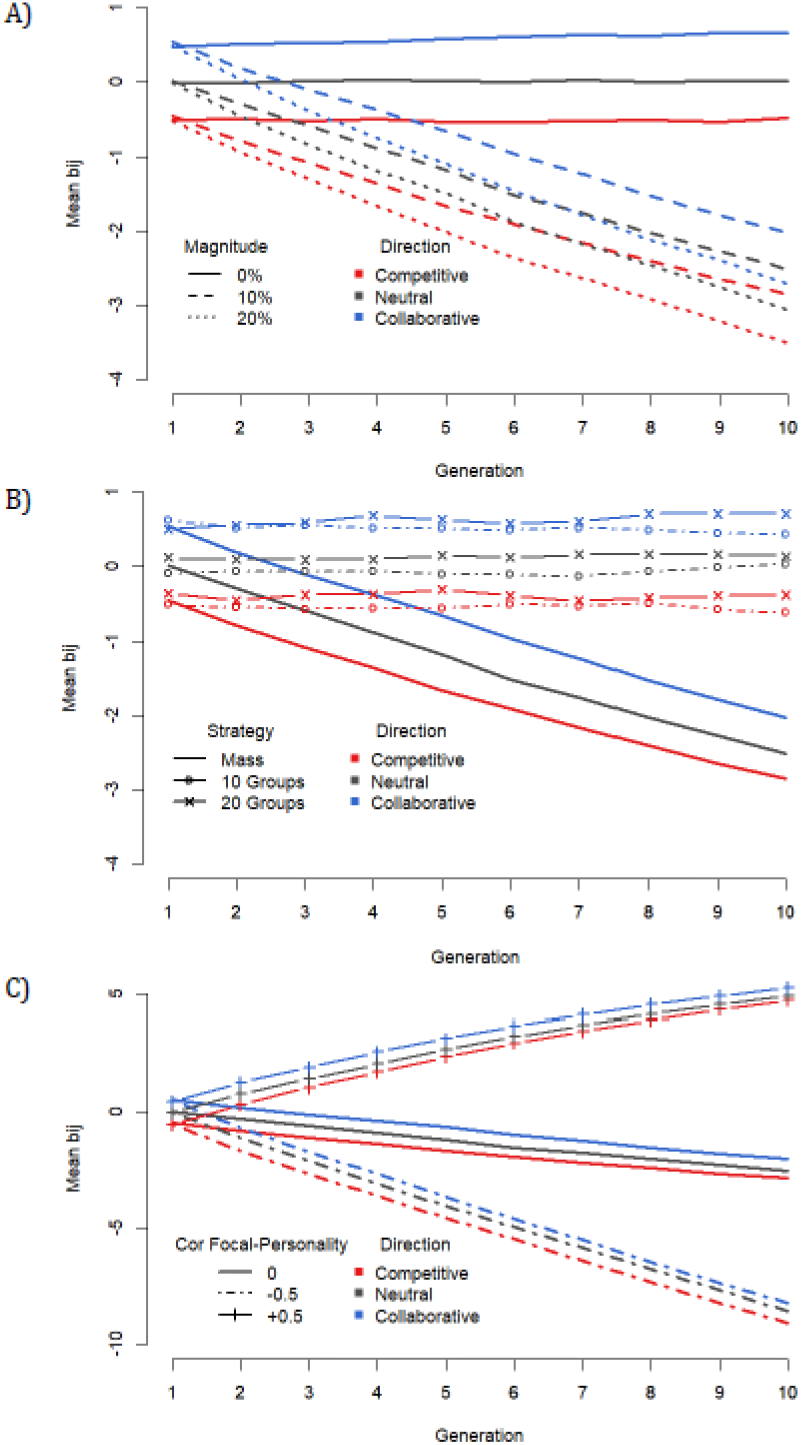
Evolution of the average b_i,j_ value over 10 generations in different scenarios: A) different magnitude of social interactions’ effects on the phenotypic variance of the focal trait; B) three selection strategies with a scenario of 10 % of the phenotypic variance of the focal trait; C) direction of the genetic and environmental correlation between focal and personality traits with a scenario of 10 % of the phenotypic variance of the focal trait.

### Selection strategy

In a scenario with SI effects (SI10) and uncorrelated focal and personality traits, we evaluated the outcomes of different selection strategies. Phenotypic mass selection consistently produced higher phenotypic and genetic means by generation 10 across all *b* scenarios, while also resulting in a substantial increase in phenotypic variance, as previously mentioned. Group selection showed reduced improvement relative to mass selection in the phenotypic and genetic means of the focal trait. However, group selection did not increase the phenotypic variance. These results align with those reported by Marjanovic *et al*. (2018), in a model in which the focal trait was also the effector trait of SIs, observed an increase in the phenotypic variance of the focal trait under mass selection, and only a modest increase under group selection. The authors suggest that, based on kin selection theory, related individuals have less variability; consequently, groups consisting of relatives have greater bij, allowing canalization (suppression of phenotypic variation) through kin selection.

Our study examined three scenarios with differential selection pressures at three levels: individuals, families, and groups of families (Wade, 1982). Thus, the selection response depends on both the relatedness among interacting individuals and the selection pressures across these multiple levels (Griffing, 1967, 1976; Bijma, Muir, and Van Arendonk, 2007; Bijma and Wade, 2008).

Mass selection emphasized individual-level selection pressure, while the two group selection strategies balanced the number of families (groups) and within-group relatedness. With the reduction in the number of families per group from 100 (10 males × 10 females) to 25 (5 males × 5 females), the full-sib groups were larger. In this setup, the reduction in the number of families led to greater genetic (and phenotypic) progress. Our empirical results agree with (Bijma, Muir, Ellen, *et al*., 2007), who showed that group selection with higher within-group relatedness yielded a higher response. Additionally, both group selection strategies (5 or 10 female/male combinations) reduced variance over the 10 simulated generations, consistent with Marjanovic *et al*. (2018).

In our study, group selection resulted in negligible or minor genetic changes in both personality traits, favoring more collaborative personalities and greater resilience to social environment effects. Thus, group selection did not change the mean b_ij_, unlike mass selection did (Fig. 8.B). To outperform, a group has to prioritize collective performance. In this sense, group selection might favor individuals with positive contribution to both, the focal trait and the social environment, thereby minimizing changes in personality traits or favoring changes in collaborative behavior rather than competitive ones. Several studies reported in breeding contexts, that mass selection increased aggression, whereas group-based selection improved performance and reduced adverse behavioral responses (Muir, 1996; Ellen *et al*., 2014; Muir *et al*., 2014).

### Trait correlations

With SI effects affecting 10% of the phenotypic variance of the focal trait, three scenarios of genetic correlations (0, -0.5, and +0.5) between the focal trait and the two personality traits, uncorrelated with each other, were assessed. The scenario with negative correlation showed greater phenotypic and genetic mean of the focal trait, beside the sign of the SIs (*b*), at generation 10. When all the traits were uncorrelated, the results were intermediate and the lowest gain was in the scenario with positive correlations. However, a dramatic difference was observed in the variance of the focal trait. In the scenario of traits negatively correlated, at generation 10, the variance has a huge increase and was greater when the SI effects were competitive (*b*= -0.5) and lowest when they were collaborative (*b*= +0.5), but in all cases the phenotypic variance grew substantially more than the base scenario. That could be explained because this scenario might promote animals more competitive, and at the same time, less resilience to social environments. Exactly the opposite happened with positively correlated traits, beside the direction of the SI effects.

Animal behavior in domestic and natural environments is a topic of interest that has been reported extensively (Hamilton, 1971; Cole and Noakes, 1980; Abbott and Dill, 1989). However, the information on the effect of the personality/behavior traits in large groups, considering the collective behavior, is scarcer and sometimes ignores the medium-long term performance and the potential for adaptation and learning capacity of the communities over time (Biro *et al*., 2016). In the case of fish, the literature describes consistent behavioral differences and coping styles between individuals over time. As an example, in rainbow trout, some individuals display proactive personalities, characterized by higher aggression and risk-taking behavior, while others exhibit reactive tendencies, marked by lower aggression and a more cautious approach to novel environments (Millot *et al*., 2014; Lallias *et al*., 2017; Best *et al*., 2023). In general, proactive individuals can be characterized by a fight reaction, active avoidance, or high sympathetic reactivity, with relatively lower plasma cortisol levels; in contrast, the less proactive show flexibility in their behavioral responses (Millot *et al*., 2014). More competitive animals might also be more susceptible to their social environment, if the hierarchical structure is determined by the competitiveness and dominance (or equilibrium) of the participants. In contrast, those who are less competitive tend to engage less in SIs and to be more resilient to their social environment (Sloman *et al*., 2000; Cañon Jones *et al*., 2010; Gesto, 2019). However, there are no explicit references to the correlation parameters between personality traits and growth traits beyond the experimental conditions. Even more, at the time of translating the differences in behavior into like-growing traits, results are unclear. Sloman *et al*. (2000) found that the most and least dominant brown trout (*Salmo trutta*) groups increased in weight over the course of the experiment. In contrast, the intermediate group did not. This resulted in significant differences in final weight between the three groups, favoring the more dominant ones. In a farmed rainbow trout (*Oncorhynchus mykiss*) population, Gesto (2019) observed that the group of more competitive fish did not outperform in usual stocking density conditions, and did not find a clear performance benefit in selecting fish of a specific coping style for fish farming.

## CONCLUSION

In scenarios where animals are raised in large groups, our study demonstrates that increased SI effects lead to greater phenotypic variability in the trait of interest. This increased variability might has significant economic implications. Under mass selection for the focal trait, the genetic means of personality traits will change and will affect the mean coefficient of personality interactions (b_ij_). The magnitude and direction of these changes depend on the correlation, both in strength and sign, between the focal and personality traits. It is important to note that even when traits are uncorrelated, mass selection on the focal trait indirectly affects personality traits through their involvement in SIs, leading to indirect directional selection on them.

Group selection, by contrast, will not increase phenotypic variability, though it yields lower genetic gains for the focal trait than mass selection. When the focal trait is the effector trait of SIs but is modulated by latent traits such as personality traits, the correlation between the focal and latent traits will influence genetic gain and, more critically, the evolution of phenotypic variance in the focal trait.

In scenarios where personality traits influence the probability and intensity of interactions, group selection achieves genetic progress without increasing phenotypic variance in the focal trait. Moreover, it will not change the genetic mean of personality traits, thereby it will not have changes in the mean of the coefficient of personality interactions (b_ij_). However, this approach will increase the rate of inbreeding per generation, an issue that can be addressed by optimizing the number of families within each group.

## ACKNOWLEDGEMENTS

This study was carried out as part of the CoBreeding project, and received support from the French government managed by the National Research Agency under the France 2030 initiative, reference ANR-22-PEAE-0003.

## AUTHOR CONTRIBUTIONS

GR conceived and designed the study, developed the R code and performed the simulations, analyzed and interpreted the results, and wrote the manuscript. BCDC was involved in the conception of the study, revised the statistical analysis, contributed for the simulation scheme, and on the manuscript. FP supervised and co-conceived the study, participated in the interpretation of the results and in the writing of the manuscript. All authors read and approved the final version of the manuscript.

## COMPETING INTERESTS

The authors declare no competing interests.

### Declaration of generative AI and AI-assisted technologies in the writing process

While preparing this work, the first author used Grammarly to suggest grammar/language corrections/clarifications to the text drafted. After using these tools, the authors reviewed and edited the content as needed and took full responsibility for the publication’s content.

## SUPPLEMENTARY TABLES

**Table S1.a.** Least squares means and group of significant differences of mean observed phenotype (P observed), in generations 1 and 10, for three scenarios of magnitude of social interaction effects (0, 10 and 20% of phenotypic variance) and three scenarios of mean direction of the social interactions 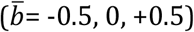.

| Generation | Trait | Scenario | Beta | LSmean | Significance group |
| --- | --- | --- | --- | --- | --- |
| 1 | $\bar{P}$ observed | 20 % | -0.5 | 0.0082 | a |
| 1 | $\bar{P}$ observed | 20 % | 0 | 0.0176 | a |
| 1 | $\bar{P}$ observed | 20 % | +0.5 | 0.0159 | a |
| 1 | $\bar{P}$ observed | 10 % | -0.5 | 0.0004 | a |
| 1 | $\bar{P}$ observed | 10 % | 0 | -0.0021 | a |
| 1 | $\bar{P}$ observed | 10 % | +0.5 | 0.0078 | a |
| 1 | $\bar{P}$ observed | 0 % | -0.5 | 0.0082 | a |
| 1 | $\bar{P}$ observed | 0 % | 0 | -0.0006 | a |
| 1 | $\bar{P}$ observed | 0 % | +0.5 | 0.0017 | a |
| 10 | $\bar{P}$ observed | 20 % | -0.5 | 8.0773 | a |
| 10 | $\bar{P}$ observed | 20 % | 0 | 7.9566 | b |
| 10 | $\bar{P}$ observed | 20 % | +0.5 | 7.8950 | b |
| 10 | $\bar{P}$ observed | 10 % | -0.5 | 7.7724 | c |
| 10 | $\bar{P}$ observed | 10 % | 0 | 7.7031 | cd |
| 10 | $\bar{P}$ observed | 10 % | +0.5 | 7.6326 | d |
| 10 | $\bar{P}$ observed | 0 % | -0.5 | 7.6412 | d |
| 10 | $\bar{P}$ observed | 0 % | 0 | 7.6647 | d |
| 10 | $\bar{P}$ observed | 0 % | +0.5 | 7.6465 | d |

**Table S1.b.** Least squares means and group of significant differences of the mean variance of the observed phenotype, in generations 1 and 10, for three scenarios of magnitude of social interaction effects (0, 10 and 20% of phenotypic variance) and three scenarios of mean direction of the social interactions 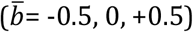.

| Generation | Trait | Scenario | Beta | LSMean | Significance group |
| --- | --- | --- | --- | --- | --- |
| 1 | $\sigma^2 P_{observed}$ | 20 % | -0.5 | 3.7190 | a |
| 1 | $\sigma^2 P_{observed}$ | 20 % | 0 | 3.0089 | c |
| 1 | $\sigma^2 P_{observed}$ | 20 % | +0.5 | 2.4322 | e |
| 1 | $\sigma^2 P_{observed}$ | 10 % | -0.5 | 3.4582 | b |
| 1 | $\sigma^2 P_{observed}$ | 10 % | 0 | 2.9684 | c |
| 1 | $\sigma^2 P_{observed}$ | 10 % | +0.5 | 2.5313 | d |
| 1 | $\sigma^2 P_{observed}$ | 0 % | -0.5 | 2.9711 | c |
| 1 | $\sigma^2 P_{observed}$ | 0 % | 0 | 3.0104 | c |
| 1 | $\sigma^2 P_{observed}$ | 0 % | +0.5 | 2.9960 | c |
| 10 | $\sigma^2 P_{observed}$ | 20 % | -0.5 | 8.3903 | a |
| 10 | $\sigma^2 P_{observed}$ | 20 % | 0 | 7.5414 | b |
| 10 | $\sigma^2 P_{observed}$ | 20 % | +0.5 | 6.8174 | c |
| 10 | $\sigma^2 P_{observed}$ | 10 % | -0.5 | 5.8037 | d |
| 10 | $\sigma^2 P_{observed}$ | 10 % | 0 | 5.3644 | e |
| 10 | $\sigma^2 P_{observed}$ | 10 % | +0.5 | 4.7714 | f |
| 10 | $\sigma^2 P_{observed}$ | 0 % | -0.5 | 2.7749 | g |
| 10 | $\sigma^2 P_{observed}$ | 0 % | 0 | 2.7742 | g |
| 10 | $\sigma^2 P_{observed}$ | 0 % | +0.5 | 2.7707 | g |

**Table S2a.**
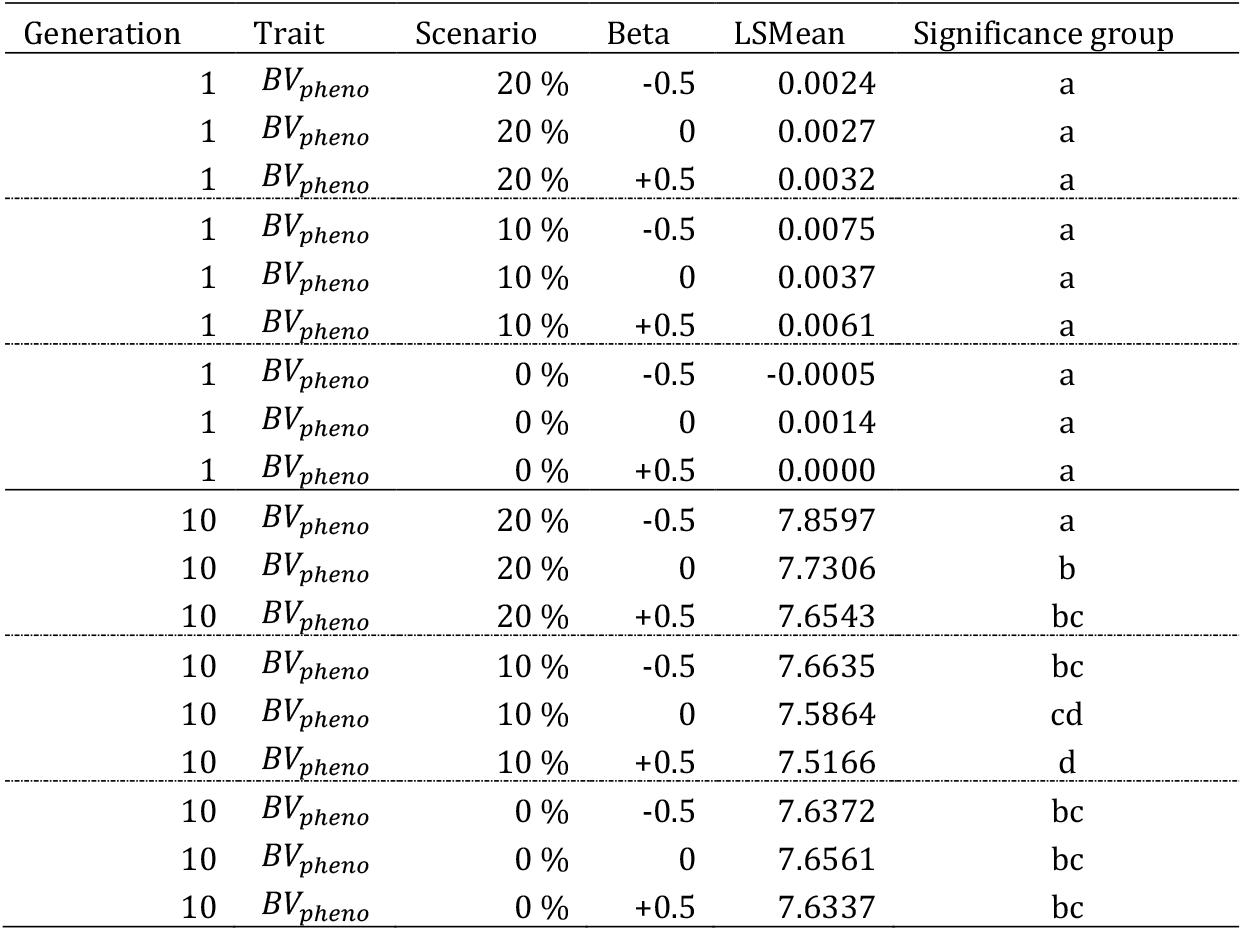
Least squares means and group of significant differences of the mean genetic value of the focal trait (BV_pheno_), in generations 1 and 10, for three scenarios of magnitude of social interaction effects (0, 10 and 20% of phenotypic variance) and three scenarios of mean direction of the social interactions 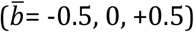.

**Table S2.b.** Least squares means and group of significant differences of the mean genetic value of personality trait (BV_P_) linked to collaborative-competitive behavior, in generations 1 and 10, for three scenarios of magnitude of social interaction effects (0, 10 and 20% of phenotypic variance) and three scenarios of mean direction of the social interactions 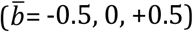.

| Generation | Trait | Scenario | Beta | LSMean | Significance group |
| --- | --- | --- | --- | --- | --- |
| 1 | $BV_P$ | 20 % | -0.5 | 0.0063 | a |
| 1 | $BV_P$ | 20 % | 0 | 0.0035 | a |
| 1 | $BV_P$ | 20 % | +0.5 | 0.0047 | a |
| 1 | $BV_P$ | 10 % | -0.5 | -0.0060 | a |
| 1 | $BV_P$ | 10 % | 0 | -0.0041 | a |
| 1 | $BV_P$ | 10 % | +0.5 | -0.0066 | a |
| 1 | $BV_P$ | 0 % | -0.5 | -0.0108 | a |
| 1 | $BV_P$ | 0 % | 0 | -0.0082 | a |
| 1 | $BV_P$ | 0 % | +0.5 | -0.0136 | a |
| 10 | $BV_P$ | 20 % | -0.5 | -2.2457 | cd |
| 10 | $BV_P$ | 20 % | 0 | -2.5147 | e |
| 10 | $BV_P$ | 20 % | +0.5 | -2.7988 | f |
| 10 | $BV_P$ | 10 % | -0.5 | -1.9612 | b |
| 10 | $BV_P$ | 10 % | 0 | -2.1550 | c |
| 10 | $BV_P$ | 10 % | +0.5 | -2.3410 | d |
| 10 | $BV_P$ | 0 % | -0.5 | -0.0331 | a |
| 10 | $BV_P$ | 0 % | 0 | -0.0050 | a |
| 10 | $BV_P$ | 0 % | +0.5 | 0.0817 | a |

**Table S2.c.**
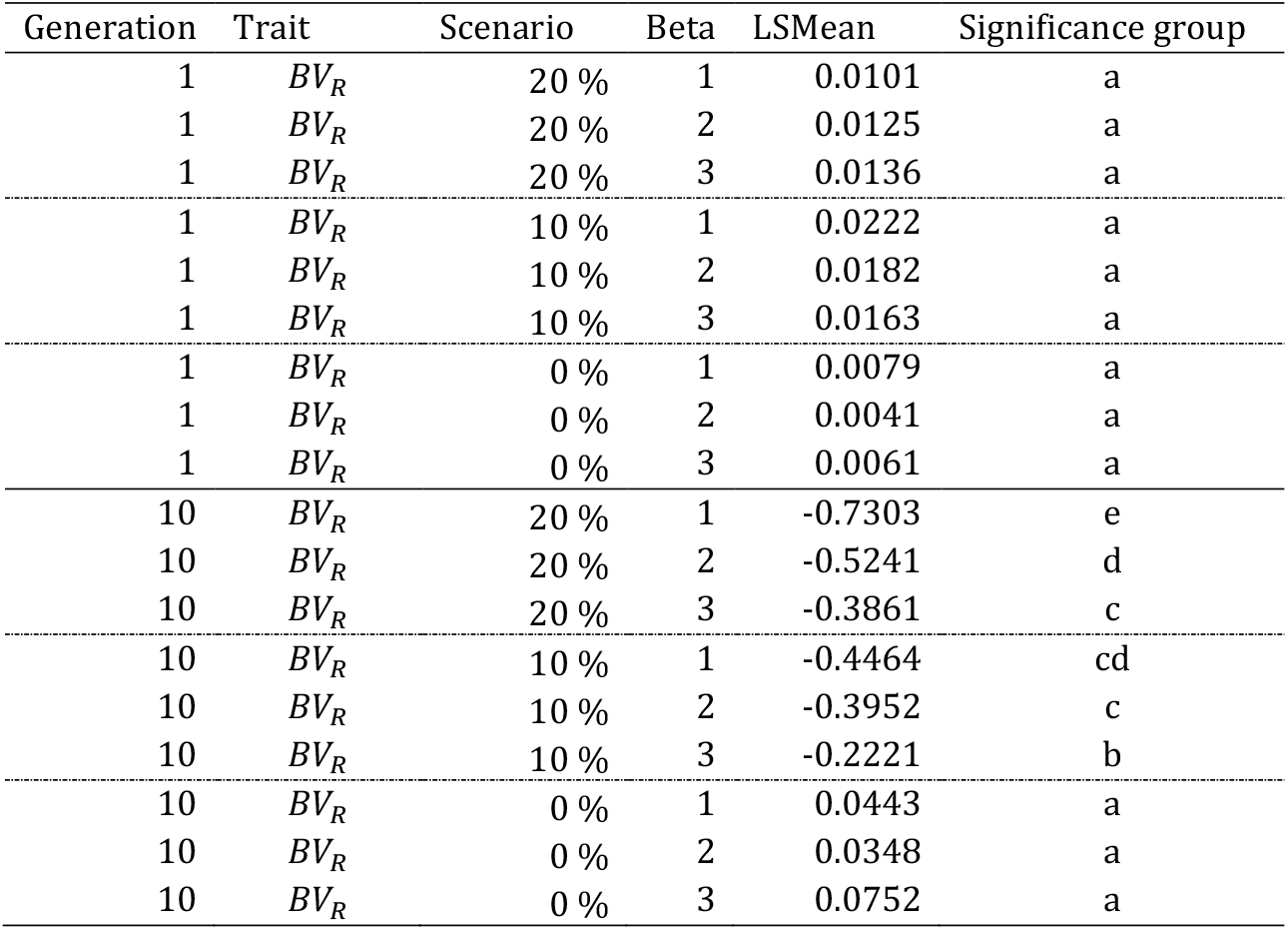
Least squares means and group of significant differences of the mean genetic value of personality trait (BV_R_) linked to resistant-susceptible to social interaction behavior, in generations 1 and 10, for three scenarios of magnitude of social interaction effects (0, 10 and 20% of phenotypic variance) and three scenarios of mean direction of the social interactions 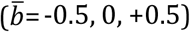.

**Table S3a.**
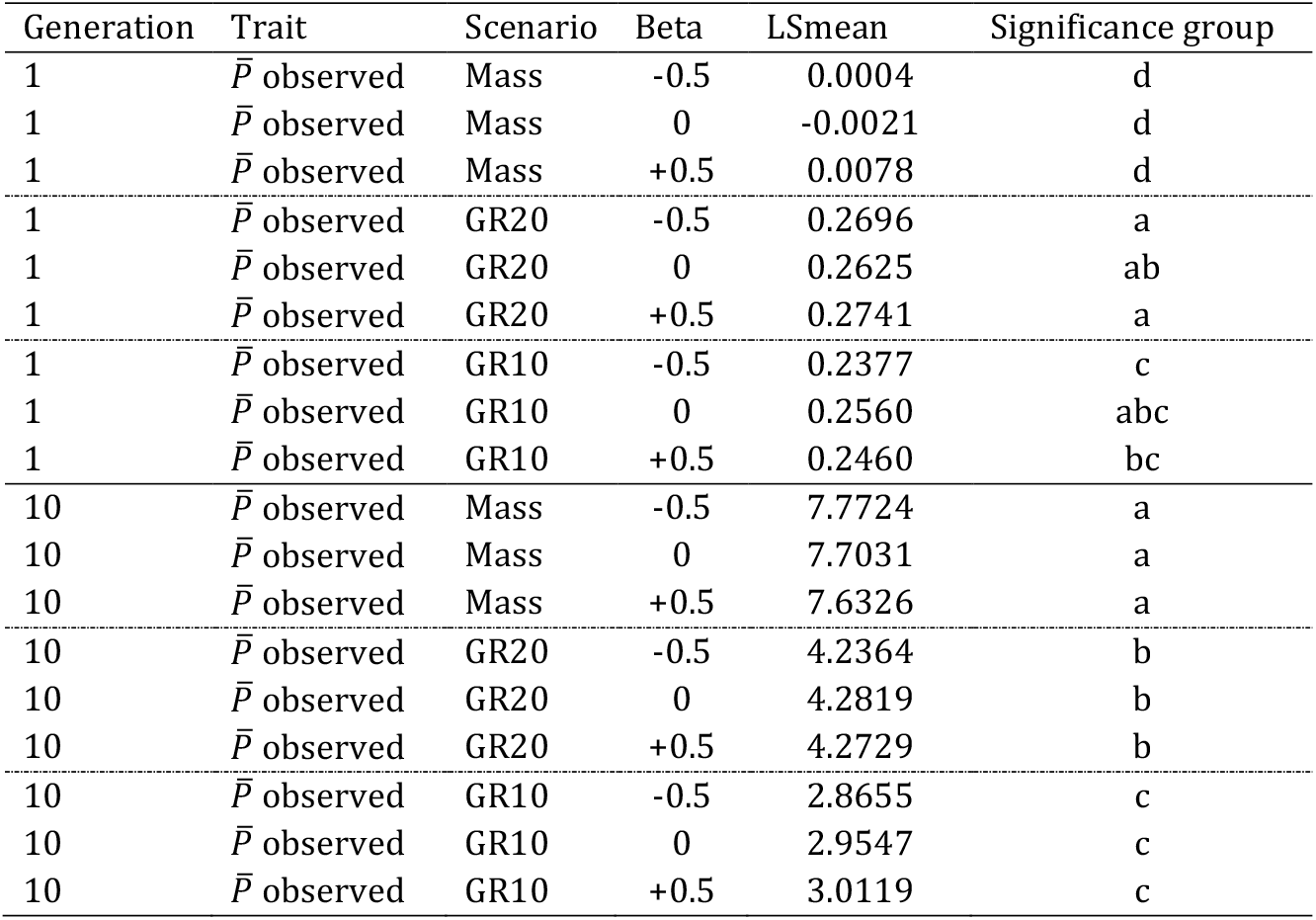
Least squares means and group of significant differences of mean observed phenotype (P observed), in generations 1 and 10, for three selection strategies: Mass; Group selection from 10 groups (GR10); and Group selection from 20 groups (GR20), and three scenarios of mean direction of the social interactions 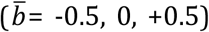. Proportion selected 10%: 200 individuals (Mass); 1 group with 200 individuals (GR10); and 2 groups with 100 individuals each (GR20).

**Table S3.b.** Least squares means and group of significant differences of the mean variance of the observed phenotype, in generations 1 and 10, for three selection strategies: Mass; Group selection from 10 families groups (GR10); and Group selection from 20 families groups (GR20) and three scenarios of mean direction of the social interactions 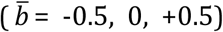. Proportion selected 10%: 200 individuals (Mass); 1 group with 200 individuals (GR10); 2 groups with 100 individuals each (GR20).

| Generation | Trait | Scenario | Beta | LSmean | Significance group |
| --- | --- | --- | --- | --- | --- |
| 1 | $\sigma^2 P_{observed}$ | Mass | -0.5 | 3.4582 | a |
| 1 | $\sigma^2 P_{observed}$ | Mass | 0 | 2.9684 | b |
| 1 | $\sigma^2 P_{observed}$ | Mass | +0.5 | 2.5313 | f |
| 1 | $\sigma^2 P_{observed}$ | GR20 | -0.5 | 2.7469 | c |
| 1 | $\sigma^2 P_{observed}$ | GR20 | 0 | 2.7174 | cd |
| 1 | $\sigma^2 P_{observed}$ | GR20 | +0.5 | 2.6484 | de |
| 1 | $\sigma^2 P_{observed}$ | GR10 | -0.5 | 2.7630 | c |
| 1 | $\sigma^2 P_{observed}$ | GR10 | 0 | 2.7188 | cd |
| 1 | $\sigma^2 P_{observed}$ | GR10 | +0.5 | 2.6323 | e |
| 10 | $\sigma^2 P_{observed}$ | Mass | -0.5 | 5.8037 | a |
| 10 | $\sigma^2 P_{observed}$ | Mass | 0 | 5.3644 | b |
| 10 | $\sigma^2 P_{observed}$ | Mass | +0.5 | 4.7714 | c |
| 10 | $\sigma^2 P_{observed}$ | GR20 | -0.5 | 2.4774 | d |
| 10 | $\sigma^2 P_{observed}$ | GR20 | 0 | 2.4521 | d |
| 10 | $\sigma^2 P_{observed}$ | GR20 | +0.5 | 2.4045 | d |
| 10 | $\sigma^2 P_{observed}$ | GR10 | -0.5 | 2.5498 | d |
| 10 | $\sigma^2 P_{observed}$ | GR10 | 0 | 2.5155 | d |
| 10 | $\sigma^2 P_{observed}$ | GR10 | +0.5 | 2.4331 | d |

**Table S4.a.** Least squares means and group of significant differences of the mean genetic value of the focal trait (BV_pheno_), in generations 1 and 10, for three selection strategies: Mass; Group selection from 10 groups (GR10); and Group selection from 20 groups (GR20) and three scenarios of mean direction of the social interactions 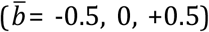. Proportion selected 10%: 200 individuals (Mass); 1 group with 200 individuals (GR10); 2 groups with 100 individuals each (GR20).

| Generation | Trait | Scenario | Beta | LSMean | Significance group |
| --- | --- | --- | --- | --- | --- |
| 1 | $BV_{pheno}$ | Mass | -0.5 | 0.0075 | a |
| 1 | $BV_{pheno}$ | Mass | 0 | 0.0037 | a |
| 1 | $BV_{pheno}$ | Mass | +0.5 | 0.0061 | a |
| 1 | $BV_{pheno}$ | GR20 | -0.5 | 0.0030 | a |
| 1 | $BV_{pheno}$ | GR20 | 0 | 0.0055 | a |
| 1 | $BV_{pheno}$ | GR20 | +0.5 | 0.0034 | a |
| 1 | $BV_{pheno}$ | GR10 | -0.5 | 0.0018 | a |
| 1 | $BV_{pheno}$ | GR10 | 0 | 0.0052 | a |
| 1 | $BV_{pheno}$ | GR10 | +0.5 | 0.0036 | a |
| 10 | $BV_{pheno}$ | Mass | -0.5 | 7.6635 | a |
| 10 | $BV_{pheno}$ | Mass | 0 | 7.5864 | a |
| 10 | $BV_{pheno}$ | Mass | +0.5 | 7.5166 | a |
| 10 | $BV_{pheno}$ | GR20 | -0.5 | 3.9650 | b |
| 10 | $BV_{pheno}$ | GR20 | 0 | 4.0105 | b |
| 10 | $BV_{pheno}$ | GR20 | +0.5 | 3.9989 | b |
| 10 | $BV_{pheno}$ | GR10 | -0.5 | 2.6144 | c |
| 10 | $BV_{pheno}$ | GR10 | 0 | 2.7086 | c |
| 10 | $BV_{pheno}$ | GR10 | +0.5 | 2.7643 | c |

**Table S4.b.** Least squares means and group of significant differences of the mean genetic value of personality trait (BV_P_) linked to collaborative-competitive behavior, in generations 1 and 10, for three selection strategies: Mass; Group selection from 10 groups (GR10); and Group selection from 20 groups (GR20) and three scenarios of mean direction of the social interactions 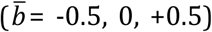. Proportion selected 10%: 200 individuals (Mass); 1 group with 200 individuals (GR10); 2 groups with 100 individuals each (GR20).

| Generation | Trait | Scenario | Beta | LSMean | Significance group |
| --- | --- | --- | --- | --- | --- |
| 1 | $BV_P$ | Mass | -0.5 | -0.0060 | a |
| 1 | $BV_P$ | Mass | 0 | -0.0041 | a |
| 1 | $BV_P$ | Mass | +0.5 | -0.0066 | a |
| 1 | $BV_P$ | GR20 | -0.5 | -0.0061 | a |
| 1 | $BV_P$ | GR20 | 0 | -0.0071 | a |
| 1 | $BV_P$ | GR20 | +0.5 | -0.0069 | a |
| 1 | $BV_P$ | GR10 | -0.5 | -0.0052 | a |
| 1 | $BV_P$ | GR10 | 0 | -0.0054 | a |
| 1 | $BV_P$ | GR10 | +0.5 | -0.0017 | a |
| 10 | $BV_P$ | Mass | -0.5 | -1.9612 | b |
| 10 | $BV_P$ | Mass | 0 | -2.1550 | bc |
| 10 | $BV_P$ | Mass | +0.5 | -2.3410 | c |
| 10 | $BV_P$ | GR20 | -0.5 | 0.1340 | a |
| 10 | $BV_P$ | GR20 | 0 | -0.0278 | a |
| 10 | $BV_P$ | GR20 | +0.5 | 0.0166 | a |
| 10 | $BV_P$ | GR10 | -0.5 | -0.0963 | a |
| 10 | $BV_P$ | GR10 | 0 | -0.1105 | a |
| 10 | $BV_P$ | GR10 | +0.5 | -0.0643 | a |

**Table S4.c.** Least squares means and group of significant differences of the mean genetic value of personality trait (BV_R_) linked to resistant-susceptible to social interaction behavior, in generations 1 and 10, for three selection strategies: Mass; Group selection from 10 groups (GR10); and Group selection from 20 groups (GR20) and three scenarios of mean direction of the social interactions 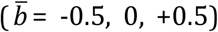. Proportion selected 10%: 200 individuals (Mass); 1 group with 200 individuals (GR10); 2 groups with 100 individuals each (GR20).

| Generation | Trait | Scenario | Beta | LSMean | Significance group |
| --- | --- | --- | --- | --- | --- |
| 1 | $BV_R$ | Mass | -0.5 | 0.0222 | a |
| 1 | $BV_R$ | Mass | 0 | 0.0182 | a |
| 1 | $BV_R$ | Mass | +0.5 | 0.0163 | a |
| 1 | $BV_R$ | GR20 | -0.5 | 0.0170 | a |
| 1 | $BV_R$ | GR20 | 0 | 0.0166 | a |
| 1 | $BV_R$ | GR20 | +0.5 | 0.0217 | a |
| 1 | $BV_R$ | GR10 | -0.5 | 0.0171 | a |
| 1 | $BV_R$ | GR10 | 0 | 0.0140 | a |
| 1 | $BV_R$ | GR10 | +0.5 | 0.0149 | a |
| 10 | $BV_R$ | Mass | -0.5 | -0.4464 | d |
| 10 | $BV_R$ | Mass | 0 | -0.3952 | cd |
| 10 | $BV_R$ | Mass | +0.5 | -0.2221 | bcd |
| 10 | $BV_R$ | GR20 | -0.5 | -0.0501 | abc |
| 10 | $BV_R$ | GR20 | 0 | 0.1361 | ab |
| 10 | $BV_R$ | GR20 | +0.5 | 0.1649 | a |
| 10 | $BV_R$ | GR10 | -0.5 | 0.0377 | ab |
| 10 | $BV_R$ | GR10 | 0 | 0.0906 | ab |
| 10 | $BV_R$ | GR10 | +0.5 | -0.0246 | abc |

**Table S4.d.**
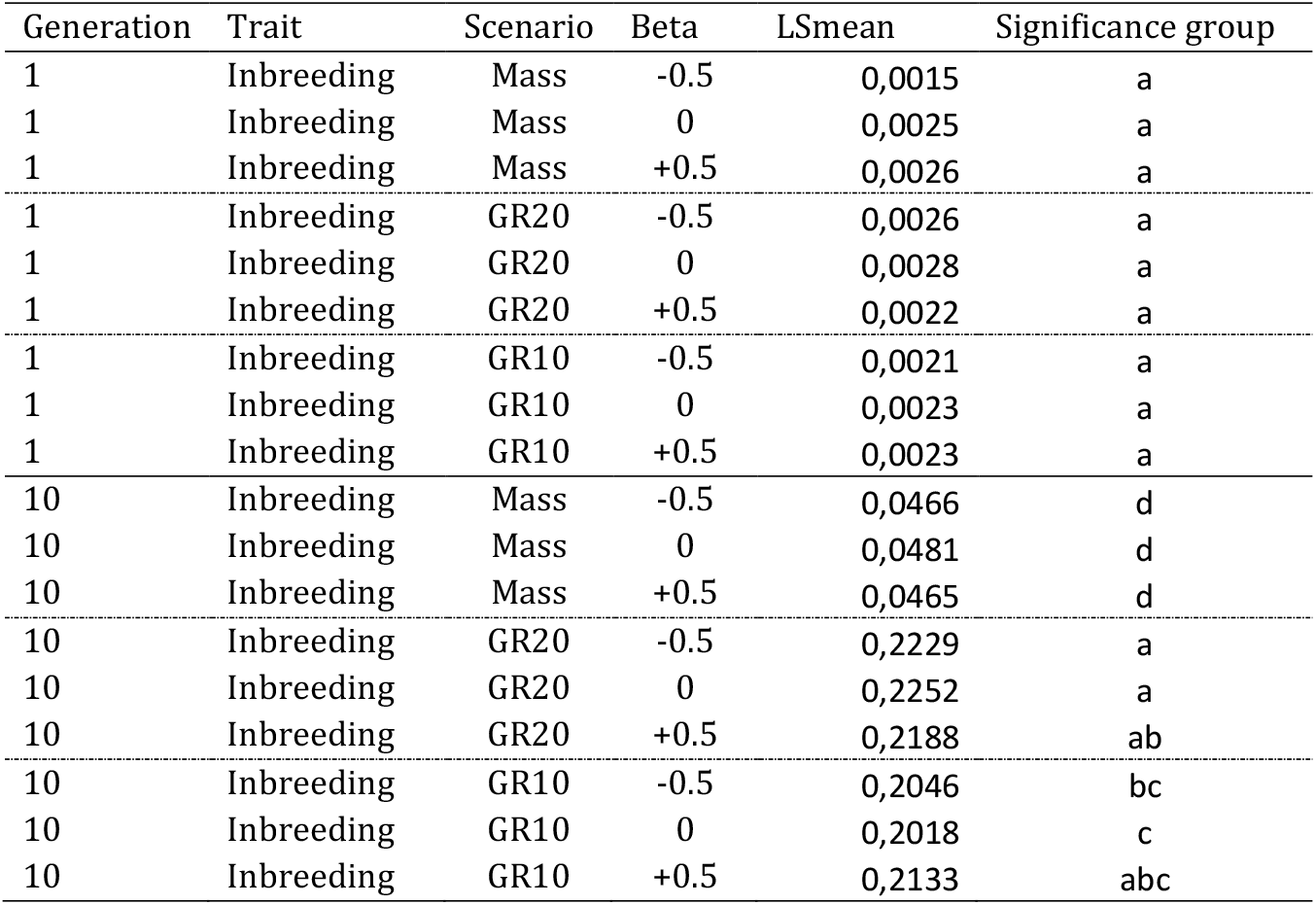
Least squares means and group of significant differences of the mean inbreeding, in generations 1 and 10, for three selection strategies: Mass; Group selection from 10 groups (GR10); and Group selection from 20 groups (GR20) and three scenarios of mean direction of the social interactions 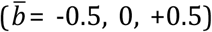. Proportion selected 10%: 200 individuals (Mass); 1 group with 200 individuals (GR10); 2 groups with 100 individuals each (GR20).

**Table S5.a.**
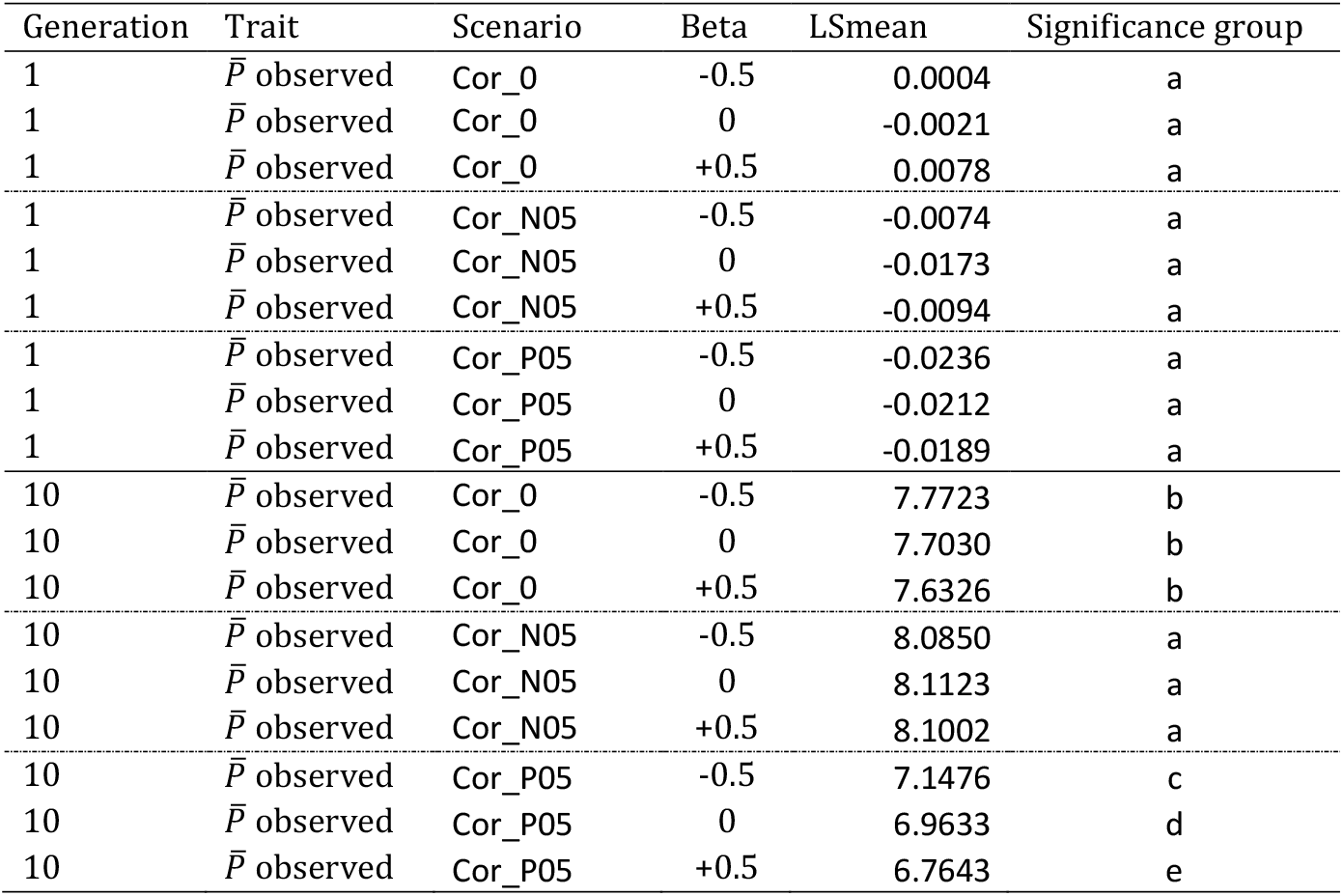
Least squares means and group of significant differences of mean observed phenotype (P observed), in generations 1 and 10, for three scenarios of correlations between the focal trait and personality traits, with a magnitude of social interaction effects of 10 % of phenotypic variance, and three scenarios of mean direction of the social interactions 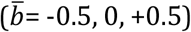. Cor_0 = Genetic and environmental correlation zero; Cor_N05 = Genetic and environmental correlation -0.5; Cor_P05 = Genetic and environmental correlation +0.5.

| Generation | Trait | Scenario | Beta | LSmean | Significance group |
| --- | --- | --- | --- | --- | --- |
| 1 | $\bar{P}$ observed | Cor_0 | -0.5 | 0.0004 | a |
| 1 | $\bar{P}$ observed | Cor_0 | 0 | -0.0021 | a |
| 1 | $\bar{P}$ observed | Cor_0 | +0.5 | 0.0078 | a |
| 1 | $\bar{P}$ observed | Cor_N05 | -0.5 | -0.0074 | a |
| 1 | $\bar{P}$ observed | Cor_N05 | 0 | -0.0173 | a |
| 1 | $\bar{P}$ observed | Cor_N05 | +0.5 | -0.0094 | a |
| 1 | $\bar{P}$ observed | Cor_P05 | -0.5 | -0.0236 | a |
| 1 | $\bar{P}$ observed | Cor_P05 | 0 | -0.0212 | a |
| 1 | $\bar{P}$ observed | Cor_P05 | +0.5 | -0.0189 | a |
| 10 | $\bar{P}$ observed | Cor_0 | -0.5 | 7.7723 | b |
| 10 | $\bar{P}$ observed | Cor_0 | 0 | 7.7030 | b |
| 10 | $\bar{P}$ observed | Cor_0 | +0.5 | 7.6326 | b |
| 10 | $\bar{P}$ observed | Cor_N05 | -0.5 | 8.0850 | a |
| 10 | $\bar{P}$ observed | Cor_N05 | 0 | 8.1123 | a |
| 10 | $\bar{P}$ observed | Cor_N05 | +0.5 | 8.1002 | a |
| 10 | $\bar{P}$ observed | Cor_P05 | -0.5 | 7.1476 | c |
| 10 | $\bar{P}$ observed | Cor_P05 | 0 | 6.9633 | d |
| 10 | $\bar{P}$ observed | Cor_P05 | +0.5 | 6.7643 | e |

**Table S5.b.** Least squares means and group of significant differences of the mean variance of the observed phenotype, in generations 1 and 10, for three scenarios of correlations between the focal trait and personality traits, with a magnitude of social interaction effects of 10 % of phenotypic variance, and three scenarios of mean direction of the social interactions 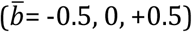. Cor_0 = Genetic and environmental correlation zero; Cor_N05 = Genetic and environmental correlation -0.5; Cor_P05 = Genetic and environmental correlation +0.5.

| Generation | Trait | Scenario | Beta | LSmean | Significance group |
| --- | --- | --- | --- | --- | --- |
| 1 | $\sigma^2 P_{observed}$ | Cor_0 | -0.5 | 3.4582 | a |
| 1 | $\sigma^2 P_{observed}$ | Cor_0 | 0 | 2.9684 | bc |
| 1 | $\sigma^2 P_{observed}$ | Cor_0 | +0.5 | 2.5313 | e |
| 1 | $\sigma^2 P_{observed}$ | Cor_N05 | -0.5 | 3.4816 | a |
| 1 | $\sigma^2 P_{observed}$ | Cor_N05 | 0 | 2.9485 | c |
| 1 | $\sigma^2 P_{observed}$ | Cor_N05 | +0.5 | 2.5263 | e |
| 1 | $\sigma^2 P_{observed}$ | Cor_P05 | -0.5 | 3.4617 | a |
| 1 | $\sigma^2 P_{observed}$ | Cor_P05 | 0 | 3.0625 | b |
| 1 | $\sigma^2 P_{observed}$ | Cor_P05 | +0.5 | 2.6812 | d |
| 10 | $\sigma^2 P_{observed}$ | Cor_0 | -0.5 | 5.8037 | d |
| 10 | $\sigma^2 P_{observed}$ | Cor_0 | 0 | 5.3644 | d |
| 10 | $\sigma^2 P_{observed}$ | Cor_0 | +0.5 | 4.7714 | e |
| 10 | $\sigma^2 P_{observed}$ | Cor_N05 | -0.5 | 17.9754 | a |
| 10 | $\sigma^2 P_{observed}$ | Cor_N05 | 0 | 16.6499 | b |
| 10 | $\sigma^2 P_{observed}$ | Cor_N05 | +0.5 | 15.9027 | c |
| 10 | $\sigma^2 P_{observed}$ | Cor_P05 | -0.5 | 0.5002 | f |
| 10 | $\sigma^2 P_{observed}$ | Cor_P05 | 0 | 0.4452 | f |
| 10 | $\sigma^2 P_{observed}$ | Cor_P05 | +0.5 | 0.3799 | f |

**Table S6a.** Least squares means and group of significant differences of the mean genetic value of the focal trait (BV_pheno_), in generations 1 and 10, for three scenarios of correlations between the focal trait and personality traits, with a magnitude of social interaction effects of 10 % of phenotypic variance, and three scenarios of mean direction of the social interactions 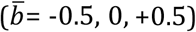. Cor_0 = Genetic and environmental correlation zero; Cor_N05 = Genetic and environmental correlation -0.5; Cor_P05 = Genetic and environmental correlation +0.5.

| Generation | Trait | Scenario | Beta | LSmean | Significance group |
| --- | --- | --- | --- | --- | --- |
| 1 | $BV_{pheno}$ | Cor_0 | -0.5 | 0,0075 | a |
| 1 | $BV_{pheno}$ | Cor_0 | 0 | 0,0037 | a |
| 1 | $BV_{pheno}$ | Cor_0 | +0.5 | 0,0061 | a |
| 1 | $BV_{pheno}$ | Cor_N05 | -0.5 | -0,0053 | a |
| 1 | $BV_{pheno}$ | Cor_N05 | 0 | -0,0100 | a |
| 1 | $BV_{pheno}$ | Cor_N05 | +0.5 | -0,0089 | a |
| 1 | $BV_{pheno}$ | Cor_P05 | -0.5 | -0,0118 | a |
| 1 | $BV_{pheno}$ | Cor_P05 | 0 | -0,0115 | a |
| 1 | $BV_{pheno}$ | Cor_P05 | +0.5 | -0,0126 | a |
| 10 | $BV_{pheno}$ | Cor_0 | -0.5 | 7,6635 | bc |
| 10 | $BV_{pheno}$ | Cor_0 | 0 | 7,5864 | c |
| 10 | $BV_{pheno}$ | Cor_0 | +0.5 | 7,5166 | c |
| 10 | $BV_{pheno}$ | Cor_N05 | -0.5 | 7,7906 | ab |
| 10 | $BV_{pheno}$ | Cor_N05 | 0 | 7,8293 | a |
| 10 | $BV_{pheno}$ | Cor_N05 | +0.5 | 7,8081 | ab |
| 10 | $BV_{pheno}$ | Cor_P05 | -0.5 | 6,9499 | d |
| 10 | $BV_{pheno}$ | Cor_P05 | 0 | 6,7676 | e |
| 10 | $BV_{pheno}$ | Cor_P05 | +0.5 | 6,5733 | f |

**Table S6.b.**
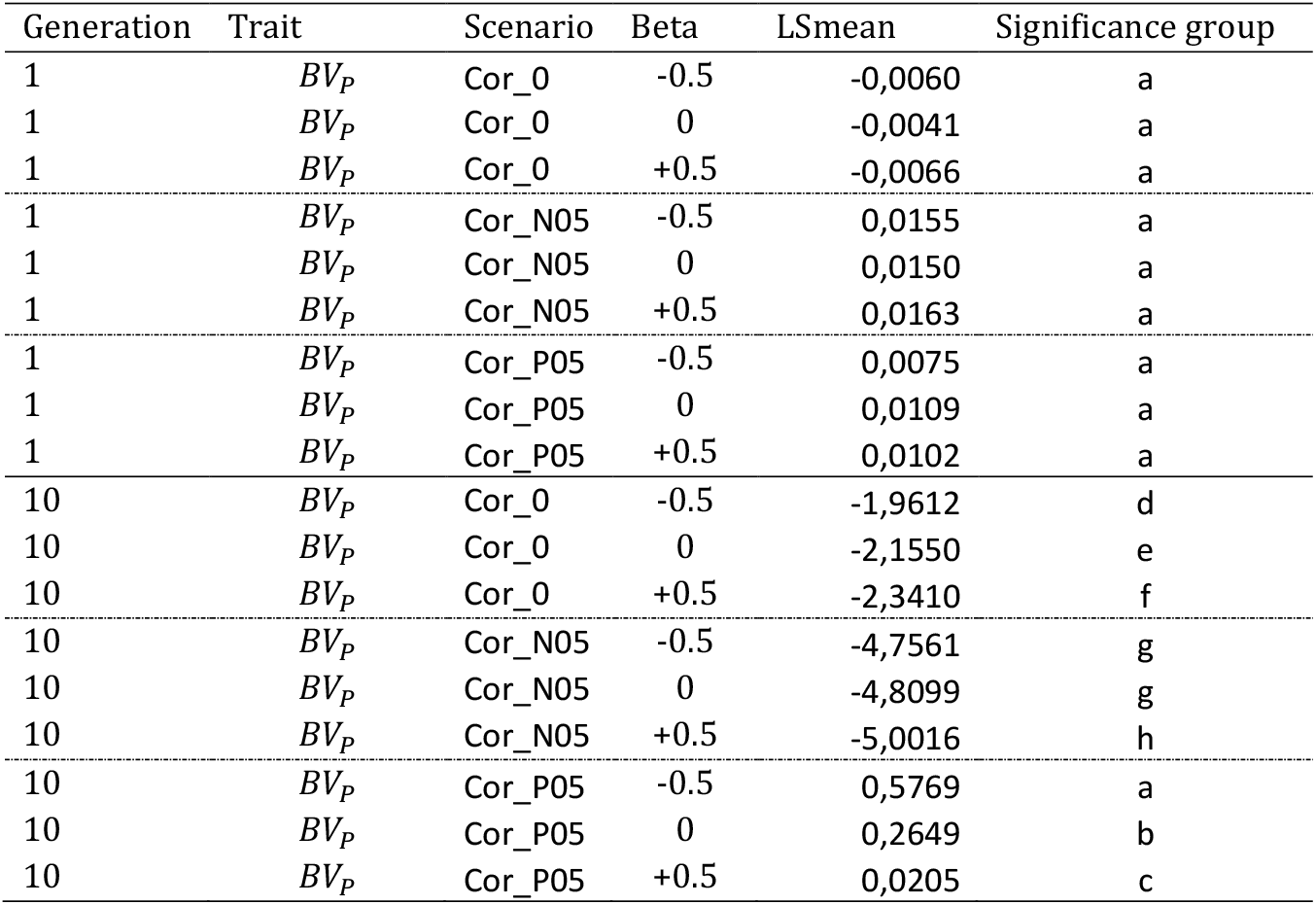
Least squares means and group of significant differences of the mean genetic value of personality trait (BV_P_) linked to collaborative-competitive behavior, in generations 1 and 10, for three scenarios of correlations between the focal trait and personality traits, with a magnitude of social interaction effects of 10 % of phenotypic variance, and three scenarios of mean direction of the social interactions 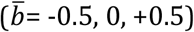. Cor_0 = Genetic and environmental correlation zero; Cor_N05 = Genetic and environmental correlation -0.5; Cor_P05 = Genetic and environmental correlation +0.5.

| Generation | Trait | Scenario | Beta | LSmean | Significance group |
| --- | --- | --- | --- | --- | --- |
| 1 | $BV_P$ | Cor_0 | -0.5 | -0,0060 | a |
| 1 | $BV_P$ | Cor_0 | 0 | -0,0041 | a |
| 1 | $BV_P$ | Cor_0 | +0.5 | -0,0066 | a |
| 1 | $BV_P$ | Cor_N05 | -0.5 | 0,0155 | a |
| 1 | $BV_P$ | Cor_N05 | 0 | 0,0150 | a |
| 1 | $BV_P$ | Cor_N05 | +0.5 | 0,0163 | a |
| 1 | $BV_P$ | Cor_P05 | -0.5 | 0,0075 | a |
| 1 | $BV_P$ | Cor_P05 | 0 | 0,0109 | a |
| 1 | $BV_P$ | Cor_P05 | +0.5 | 0,0102 | a |
| 10 | $BV_P$ | Cor_0 | -0.5 | -1,9612 | d |
| 10 | $BV_P$ | Cor_0 | 0 | -2,1550 | e |
| 10 | $BV_P$ | Cor_0 | +0.5 | -2,3410 | f |
| 10 | $BV_P$ | Cor_N05 | -0.5 | -4,7561 | g |
| 10 | $BV_P$ | Cor_N05 | 0 | -4,8099 | g |
| 10 | $BV_P$ | Cor_N05 | +0.5 | -5,0016 | h |
| 10 | $BV_P$ | Cor_P05 | -0.5 | 0,5769 | a |
| 10 | $BV_P$ | Cor_P05 | 0 | 0,2649 | b |
| 10 | $BV_P$ | Cor_P05 | +0.5 | 0,0205 | c |

**Table S6.c.** Least squares means and group of significant differences of the mean genetic value of personality trait (BV_R_) linked to resistant-susceptible to social interaction behavior, in generations 1 and 10, for three scenarios of correlations between the focal trait and personality traits, with a magnitude of social interaction effects of 10 % of phenotypic variance, and three scenarios of mean direction of the social interactions 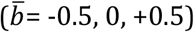. Cor_0 = Genetic and environmental correlation zero; Cor_N05 = Genetic and environmental correlation -0.5; Cor_P05 = Genetic and environmental correlation +0.5.

| Generation | Trait | Scenario | Beta | LSmean | Significance group |
| --- | --- | --- | --- | --- | --- |
| 1 | $BV_R$ | Cor_0 | -0.5 | 0,0222 | a |
| 1 | $BV_R$ | Cor_0 | 0 | 0,0182 | a |
| 1 | $BV_R$ | Cor_0 | +0.5 | 0,0163 | a |
| 1 | $BV_R$ | Cor_N05 | -0.5 | 0,0029 | a |
| 1 | $BV_R$ | Cor_N05 | 0 | 0,0068 | a |
| 1 | $BV_R$ | Cor_N05 | +0.5 | 0,0042 | a |
| 1 | $BV_R$ | Cor_P05 | -0.5 | -0,0090 | a |
| 1 | $BV_R$ | Cor_P05 | 0 | -0,0133 | a |
| 1 | $BV_R$ | Cor_P05 | +0.5 | -0,0108 | a |
| 10 | $BV_R$ | Cor_0 | -0.5 | -0,4464 | c |
| 10 | $BV_R$ | Cor_0 | 0 | -0,3952 | bc |
| 10 | $BV_R$ | Cor_0 | +0.5 | -0,2221 | b |
| 10 | $BV_R$ | Cor_N05 | -0.5 | -3,8037 | d |
| 10 | $BV_R$ | Cor_N05 | 0 | -3,7398 | d |
| 10 | $BV_R$ | Cor_N05 | +0.5 | -3,7227 | d |
| 10 | $BV_R$ | Cor_P05 | -0.5 | 4,6297 | a |
| 10 | $BV_R$ | Cor_P05 | 0 | 4,7082 | a |
| 10 | $BV_R$ | Cor_P05 | +0.5 | 4,7699 | a |

